# Opposing Modulation of EEG Aperiodic Component by Ketamine and Thiopental: Implications for the Noninvasive Assessment of Cortical E/I Balance in Humans

**DOI:** 10.1101/2025.05.12.653580

**Authors:** Jose A. Cortes-Briones, Juan Urrutia-Gandolfo, Pablo A. Estevez, Arnab Sengupta, Matthew Basilico, Patrick D. Skosnik, Diane Limoncelli, Deepak C. D’Souza, Ismene L. Petrakis, John H. Krystal

## Abstract

The balance between excitatory and inhibitory (E/I) activity is critical for brain function, and its disruption is implicated in neuropsychiatric disorders. Electrophysiological signals can be decomposed into periodic (oscillatory) and aperiodic components. In the power spectrum, the periodic component appears as narrowband peaks, while the aperiodic component underlies its characteristic 1⁄*f^x^* power-law decay. Computational models predict that shifts in E/I balance alter the exponent *χ* in specific directions. In a randomized, double-blind, placebo-controlled, within-subject study, healthy volunteers received subanesthetic doses of ketamine and thiopental during an EEG oddball task. These drugs have opposite effects on E/I balance but comparable sedative profiles. Ketamine reduced the PSD exponent, while thiopental increased it, consistent with computational predictions. Changes in the exponent were associated with subjective and cognitive effects. These findings suggest that the PSD exponent has potential as a noninvasive EEG biomarker sensitive to transient shifts in cortical E/I balance.

## Introduction

From the cellular to the global scale, brain circuits maintain a balance between excitatory and inhibitory processes^1^. Both aberrant increases and decreases in the ratio of excitation to inhibition (E/I) are associated with reduced circuit function^2^ following an inverted-U relationship^3–6^. Alterations in E/I balance also characterize a wide range of brain disorders, including epilepsy, autism, and schizophrenia^7–11^, where they are associated with impairments in cognitive function. Thus, understanding the biology of disorder- and drug-induced alterations in E/I balance could shed light on the neurobiology of psychiatric disorders and their treatment.

In the frequency domain, the power spectral density (PSD) of global electrophysiological brain signals such as local field potentials (LFPs), electroencephalography (EEG), and magnetoencephalography (MEG), decays with frequency (*f*) following a power law of the form 1/*f^x^* (*χ* > 0)^9, 12, 13^. Evidence from in silico, animal, and human studies suggests that this power law arises from an aperiodic, non-oscillatory, broadband component of neural field signals that emerges from the sum of the globally uncoordinated and mainly stochastic activity of large neural populations^14^. In addition, brain signals have a periodic component which corresponds to the well-known neural oscillations that emerge from recurrent cycles of coordinated activation and deactivation of large populations of neurons^14^. This component appears as narrowband peaks in the power spectrum^12, 13^, and is critically involved in brain functions such as perception, attention, and working memory^15^. Alterations in this component are associated with mental illness, neurological conditions, and drug intoxication^16, 17^. In contrast, the role of the aperiodic component remains poorly understood. Although it emerges from stochastic processes, it is modulated by task demands^18, 19^, and thus cannot be easily equated with noise in the traditional sense of task-unrelated activity.

Pharmacological studies in humans with drugs that transiently shift the balance between excitation and inhibition have demonstrated that shifts in E/I balance have significant effects on the periodic component (e.g., through event-related potentials and the auditory steady-state response)^20–23^, and that these effects correlate with the cognitive and perceptual effects of the drugs. In contrast, the effects that changes in E/I balance have on the aperiodic component and their relationship with behavior have received little attention in human studies. Studies have been conducted on humans under general anesthesia, which precludes the study of the subjective effects associated with controlled shifts in the PSD exponent and, more importantly, are affected by the confounding of loss of consciousness, which has profound effects on the exponent^24–26^. In these studies, propofol, a drug that increased γ-aminobutyric acid type A receptor (GABAA-R) mediated circuit inhibition, has been shown to increase the PSD exponent, while ketamine, an N-methyl-D-aspartate glutamate receptor (NMDA-R) antagonist with disinhibitory effects at low doses, has been shown to decrease it^27, 28^. Recently, the reanalysis of data from two small MEG pharmacological studies in non-anesthetized humans showed that reducing circuit inhibition decreased the PSD exponent compared to placebo (n=19), and that increasing inhibition increased the PSD exponent compared to placebo (n=15)^26^. No direct, between-study comparison of active drug conditions was conducted, nor was the relationship between changes in E/I balance and subjective effects of the drugs reported.

These findings are consistent with evidence from computer modeling and animal studies showing that changes in E/I balance are inversely correlated with the PSD exponent *χ*, particularly between 30-50Hz ^29^; i.e., the higher the E/I ratio, the smaller the *χ* (the flatter the PSD curve). However, simulations have shown that factors other than E/I ratio also have a significant influence on the PSD exponent, and that under certain conditions, the proposed relationship may not hold^14^. For example, a study in rodents failed to reduce the PSD exponent after chemogenetically reducing GABA_A_-R activity^27, 30^. Thus, to isolate the effects that changes in E/I balance have on the aperiodic component, it is essential to maintain other factors influencing the PSD exponent as constant as possible across compared conditions. These challenges and those associated with studying anesthetized subjects, could be significantly addressed with a within-subject design administering subanesthetic doses of drugs with opposite effects on E/I balance but similar behavioral profiles. To the best of our knowledge, this is the first study to implement this approach.

In this study, we addressed the confounding effects of interindividual variability and alertness using a randomized, double-blind, placebo-controlled, within-subject design, using an EEG oddball task to examine the effects of acute, drug-induced changes in E/I ratio on the aperiodic component of brain signals and their relationship with subjective effects and cognitive function. For this purpose, we administered subanesthetic doses of ketamine and thiopental, sedative drugs with opposite effects on E/I ratio but comparable behavioral profiles, to a large sample of healthy participants. Ketamine is a dissociative anesthetic^31^ and a non-competitive antagonist of the NMDA-R. Ketamine blocks NMDA-R-mediated excitation^32^, and reduces glutamate release^33^, particularly at anesthetic doses. However, at subanesthetic doses, ketamine produces a paradoxical cortical disinhibition by preferentially reducing NMDA-R-mediated excitatory input to inhibitory interneurons^34–36^, leading to increased glutamate release^33^ and increased high frequency pyramidal cell firing, albeit in a chaotic pattern associated with a suppression of bursting activity ^37, 38^. The effects of ketamine on cortical microcircuits contrast with those of another sedating medication, thiopental; a barbiturate positive allosteric modulator of GABA_A_-Rs^20, 39^. Thiopental reduces cortical excitation^40^ and decreases glutamate release^41^. These drugs were selected for having opposite effects on E/I ratio at subanesthetic doses and comparable sedative effects^20, 42^ thus, alterations in EEG measures would not be due to asymmetric changes in arousal^43^. We hypothesized that the increase in E/I-ratio induced by ketamine would reduce the PSD exponent, while the reduction in E/I ratio induced by thiopental would increase it. Furthermore, we hypothesized that the magnitude of the changes in E/I ratio would be correlated with the subjective effects and cognitive impairment of the drugs as measured by the Clinician-Administered Dissociative States Scale (CADSS)^44^ and Hopkins Verbal Learning Test (HVLT).

To better understand the relevance of the aperiodic component in capturing E/I ratio changes in the brain, we conducted data-driven analyses to assess the relative importance of the PSD exponent for identifying altered E/I states (drug conditions) against a large set of commonly used EEG measurements. We hypothesized that the PSD exponent would be the most informative biomarker for distinguishing drugs with opposite effects on E/I ratio, suggesting that changes in the aperiodic component may serve as a sensitive index of aberrant shifts in E/I balance, especially in within subject designs.

## Results

### Behavioral measures

The analyses revealed a significant main effect of drug condition on the CADSS total score (ATS=222.664, *p*<0.001). Post hoc comparisons indicated significantly higher scores in the ketamine condition compared to both placebo and thiopental (all *p*_Adj_ <0.001), and higher scores in the thiopental condition compared to placebo (*p*_Adj_<0.001). For the HVLT average score, there was a significant main effect of drug condition (ATS=44.599, p<0.001). Post hoc comparisons showed significantly lower scores under ketamine compared to both placebo and thiopental (all *p*_Adj_ <0.001), and lower scores under thiopental compared to placebo (*p*_Adj_=0.001) (see Table 1 and Figure 1).

**Figure 1.**
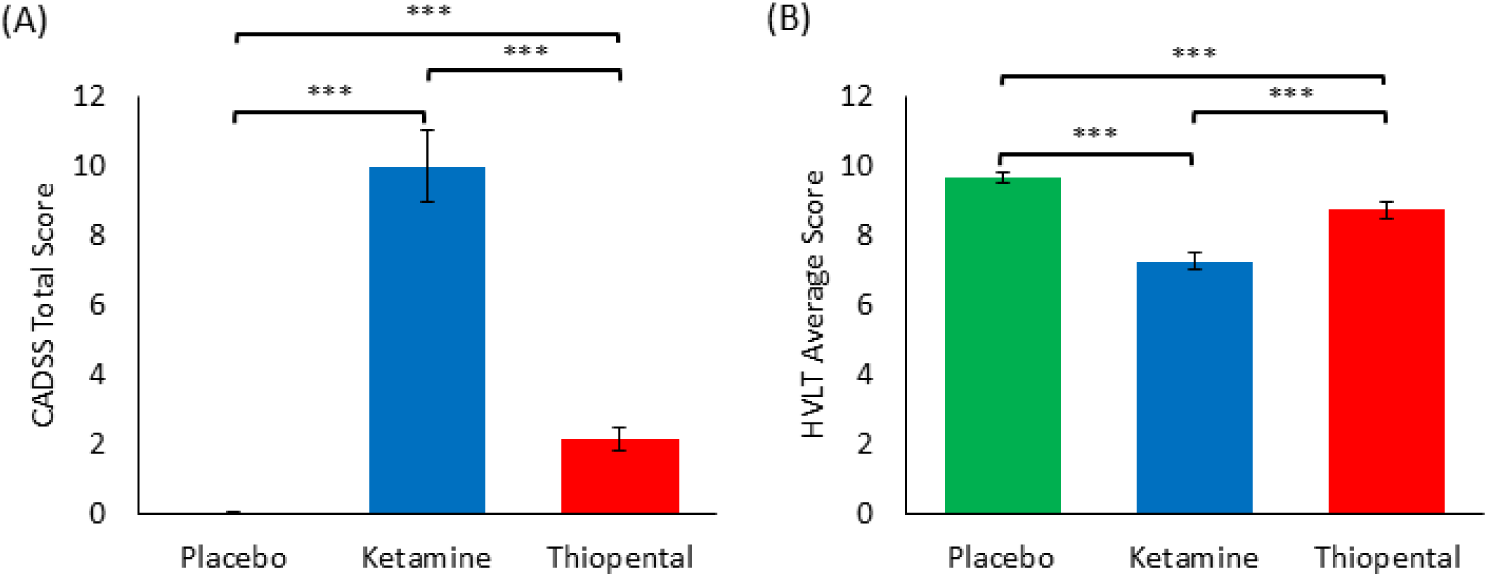
Behavioral effects across drug conditions. (A) CADSS total scores across placebo, ketamine, and thiopental conditions. (B) HVLT average scores across the same conditions. Horizontal lines and asterisks indicate significant pairwise differences (***p<0.001, Holm-Bonferroni corrected). Error bars represent the standard error of the mean (SEM).

**Table 1.** Behavioral effects across drug conditions.

| Drug Condition | CADSS Total | HVLt Average |
| --- | --- | --- |
| Placebo | 0.041 (0.200) | 9.699 (1.206) |
| Ketamine | 10.013 (8.813) | 7.280 (2.021) |
| Thiopental | 2.150 (2.543) | 8.756 (1.902) |

Means and standard deviations (SD) are reported for subjective effects (CADSS Total) and cognitive performance (HVLt Average) under each drug condition. Standard deviations are shown in parentheses.

### EEG measures

#### Power spectral density

For illustrative purposes, the average EEG power spectral density (PSD) across subjects and electrodes is shown for the post-infusion placebo, ketamine, and thiopental conditions, as well as for the pre-infusion period (Figure 2).

**Figure 2.**
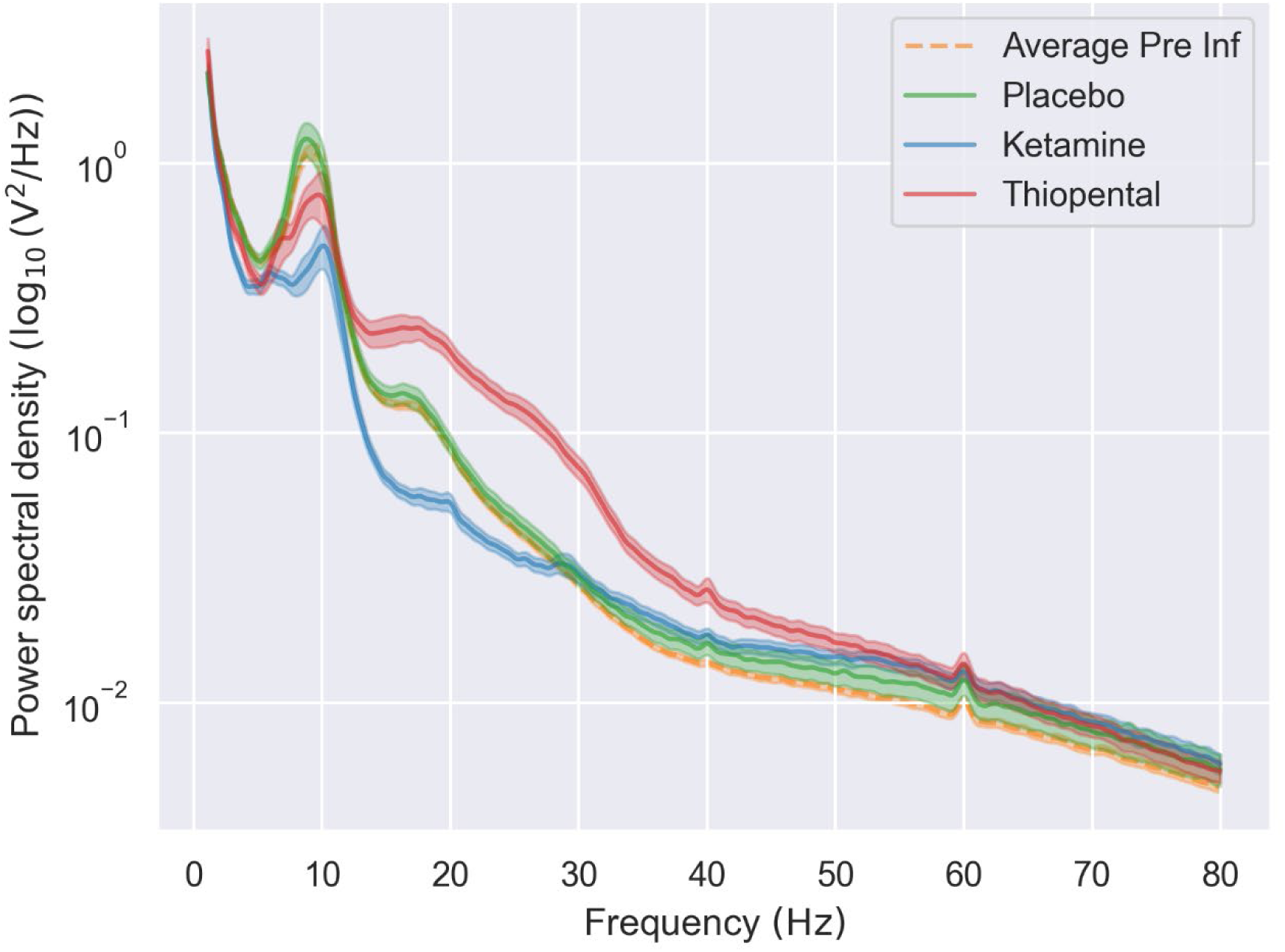
EEG power spectral density curves across drug conditions. Power spectral density (PSD) curves (log₁₀-transformed) averaged across subjects and electrodes are shown for the post-infusion placebo (green), ketamine (blue), thiopental (red), and pre-infusion placebo (dashed orange) conditions. PSDs were computed from the standard stimulus trials during the oddball task. Error shading represents the standard error of the mean.

#### Hypothesis-driven approach: Differences in PSD exponent across drug conditions

There was a main effect of drug on the absolute and the relative (change from pre-infusion to post-infusion) values of the PSD exponent during the pre-stimulation (baseline) and stimulation periods of the EEG task (*p_Adj_*<0.001, adjusted for 4 comparisons). Pairwise comparisons revealed that ketamine reduced, and thiopental increased the PSD exponent compared to placebo in all cases (*p_Adj_*<0.001, adjusted for 12 comparisons). For details, see Tables 2 and 3.

**Table 2.** Drug Effects on the PSD exponent across time windows.

| PSD exponent | Time window | Omnibus drug effect<br>(Wald $\chi^2$, $df=2$) | $p$-value<br>(Corrected) | Drug comparison | $p$-value<br>(corrected) |
| --- | --- | --- | --- | --- | --- |
| Absolute<br>(post-infusion) | Pre-stimulation | 306.045 | <0.0001 | Ketamine<Placebo | <0.0001 |
|  |  |  |  | Ketamine<Thiopental | <0.0001 |
|  |  |  |  | Thiopental>Placebo | <0.0001 |
|  | Stimulation | 389.323 | <0.0001 | Ketamine<Placebo | <0.0001 |
|  |  |  |  | Ketamine<Thiopental | <0.0001 |
|  |  |  |  | Thiopental>Placebo | <0.0001 |
| Relative<br>(change from<br>Pre-infusion) | Pre-stimulation | 216.366 | <0.0001 | Ketamine<Placebo | <0.0001 |
|  |  |  |  | Ketamine<Thiopental | <0.0001 |
|  |  |  |  | Thiopental>Placebo | <0.0001 |
|  | Stimulation | 325.774 | <0.0001 | Ketamine<Placebo | <0.0001 |
|  |  |  |  | Ketamine<Thiopental | <0.0001 |
|  |  |  |  | Thiopental>Placebo | <0.0001 |

**Table 3.** Absolute and relative PSD exponent values across drug conditions and time windows.

| Drug condition | Mean PSD exponent (95% CI) |  |  |  |
| --- | --- | --- | --- | --- |
|  | Absolute (post-infusion) |  | Relative (change from pre-infusion) |  |
|  | Pre-stimulation | Stimulation | Pre-stimulation | Stimulation |
| Placebo | 2.170 (2.025, 2.315) | 2.174 (2.013, 2.335) | -0.096 (-0.187, -0.005) | -0.076 (-0.165, 0.013) |
| Ketamine | 1.483 (1.342, 1.625) | 1.386 (1.240, 1.533) | -0.758 (-0.864, -0.653) | -0.873 (-0.973, -0.772) |
| Thiopental | 2.839 (2.673, 3.005) | 2.871 (2.699, 3.044) | 0.607 (0.461, 0.753) | 0.686 (0.548, 0.824) |

Marginal means and 95% Wald confidence intervals (CI) are reported for the PSD exponent, estimated using generalized estimating equations (GEE). Results are shown separately for absolute (post-infusion) and relative (change from pre-infusion) values across pre-stimulation and stimulation time windows.

For illustrative purposes, we show the relative location of the subjects across drug conditions in a 3-dimensional space, with the PSD exponent on the X-axis and the change from pre-infusion in the PSD exponent on the Y-axis (Figure 3).

**Figure 3.**
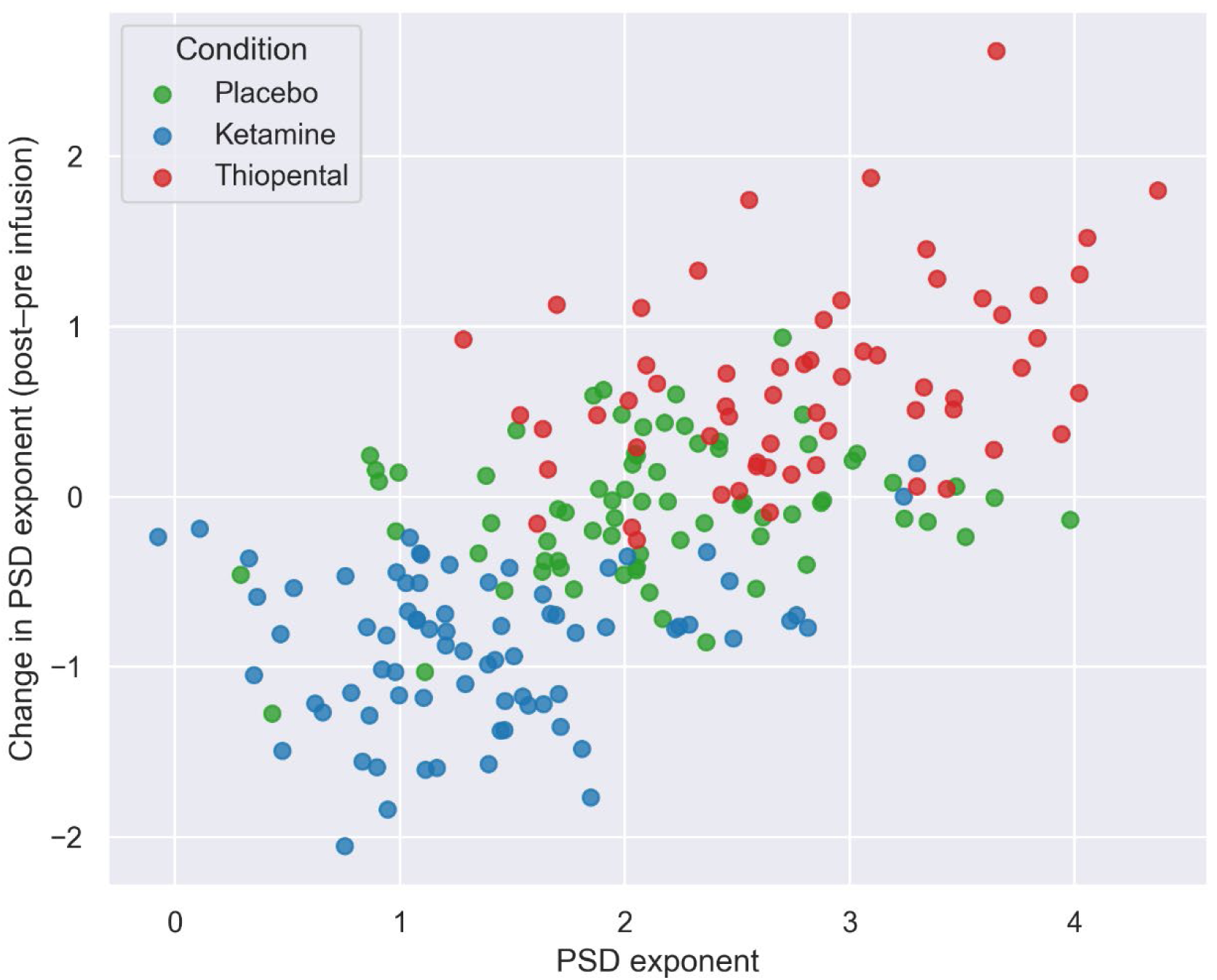
PSD exponent values per subject across drug conditions. Each point represents an individual subject. The X-axis shows the PSD exponent during post-infusion, while the Y-axis shows the change in PSD exponent from pre- to post-infusion. PSD exponents were calculated during the stimulation interval of the standard trials of the oddball task. Colors indicate drug condition (placebo, ketamine, thiopental).

##### Relationship between PSD exponent and behavior

The analyses showed significant positive associations between the PSD exponent and both the CADSS total (log_10_-transformed; Figure 4A) and HVLT average scores (Figure 4B), for both absolute (post-infusion) and relative (change from pre-infusion) PSD exponent estimates, across both pre-stimulation and stimulation periods of the oddball task (Table 4).

**Figure 4.**
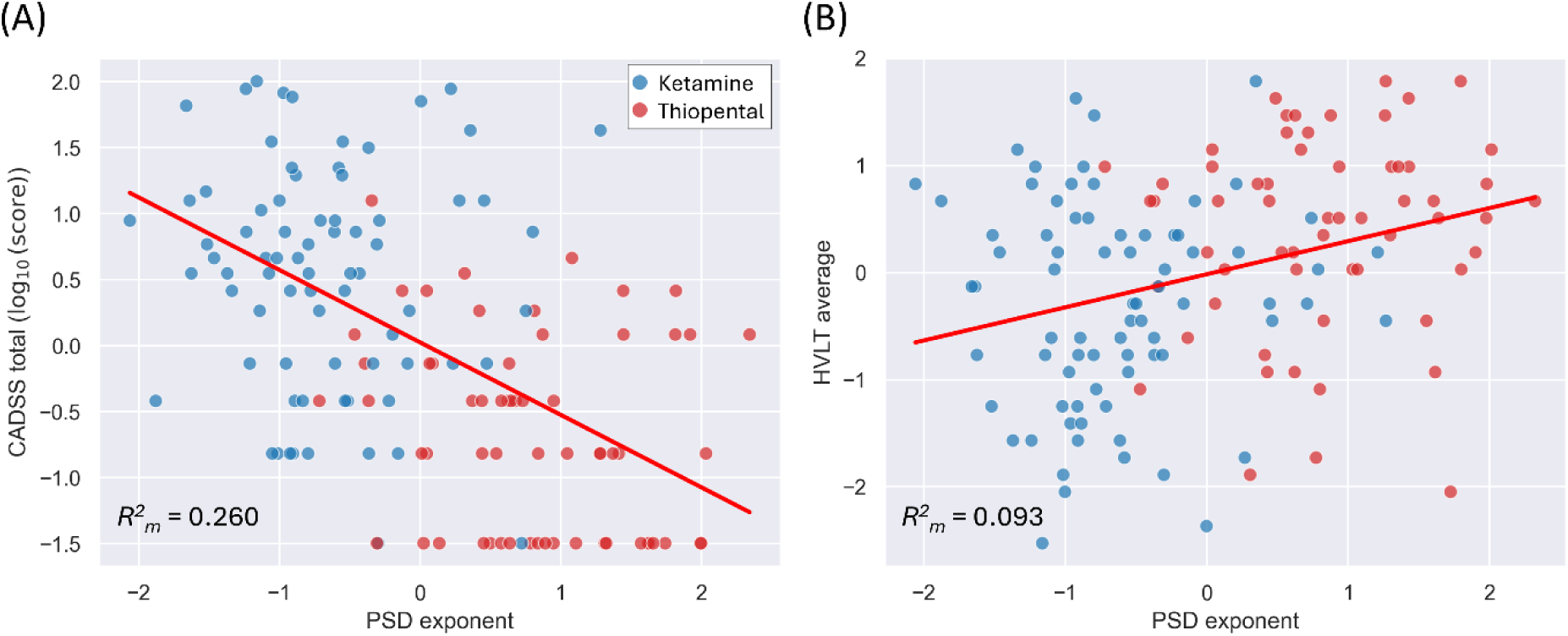
Relationship between PSD exponent and behavioral outcomes across active drug conditions. (A) Each dot represents an individual subject’s standardized (z-score) CADSS total score (log₁₀-transformed) as a function of the standardized PSD exponent during the stimulation period of standard trials. (B) Same as in (A), but with HVLT average scores. The red line indicates the GEE-based longitudinal linear regression fit across pooled data from the ketamine and thiopental conditions. Marginal R² values (R²ₘ) are shown in the bottom-left corner of each plot.

**Table 4.**
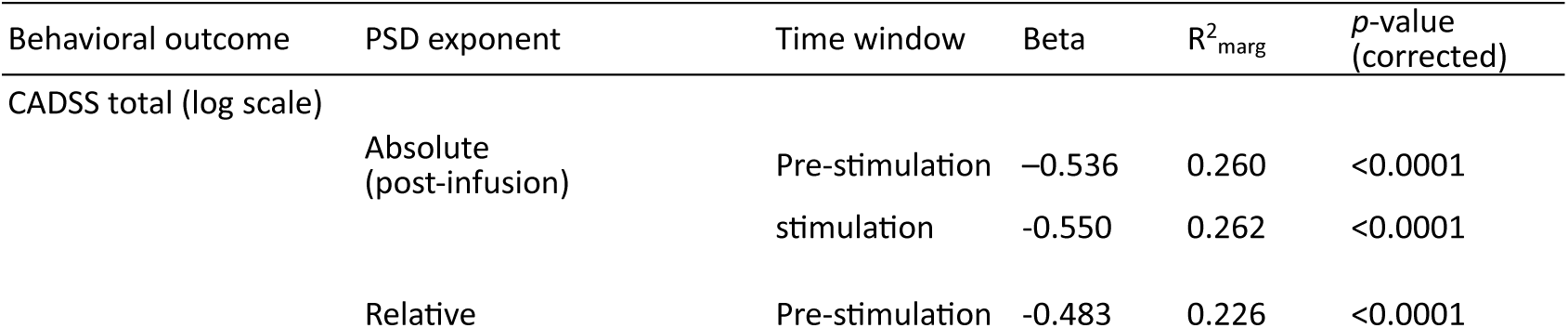
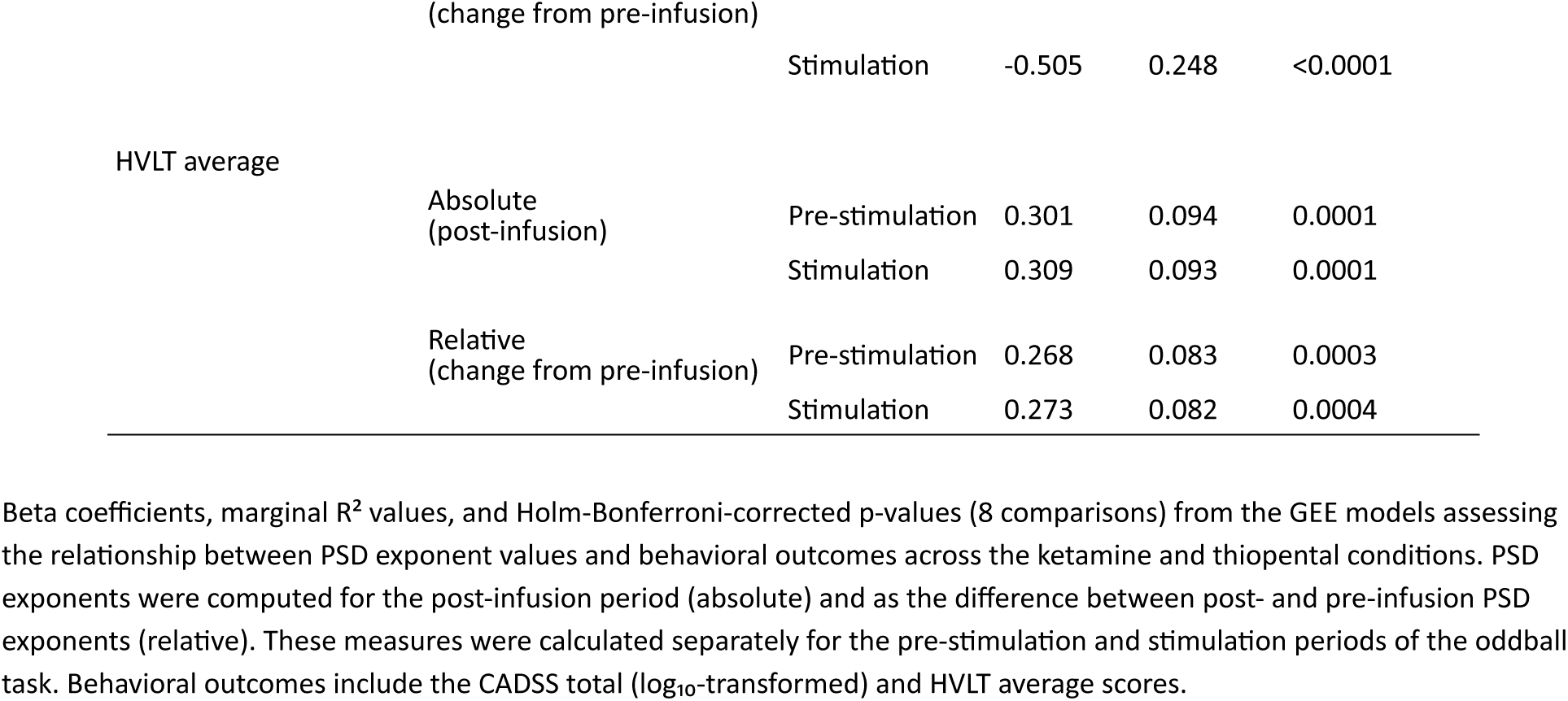
Relationship between PSD exponent values and behavioral outcomes.

#### Data-driven approach: Identification of informative EEG biomarkers for differentiating drug conditions

The EEG biomarker categories identified as the most informative for distinguishing among drug conditions are presented in Table 5. The full lists of selected biomarker categories, including those that did not meet the 50% cross-validation selection frequency threshold, are presented in Supplementary Tables S6-S9, grouped by drug comparison and selection size restriction parameter alpha. The list of biomarkers within each category is provided in Supplementary Table S1. The full lists of the selected individual EEG biomarkers (i.e., not grouped under categories) are presented in Supplementary Tables S2-S5. Classification performance metrics for each drug condition comparison and level of alpha are provided in Supplementary Tables S10.1-S10.4.

**Table 5.** Most informative EEG biomarker categories for drug condition discrimination.

| Drug Comparison | Number of Selections (Folds) | PSD Exponent (Aperiodic) | Delta Power 1<sup>st</sup> Peak (Periodic) | Theta Power (Total) | Beta Power (Total) |
| --- | --- | --- | --- | --- | --- |
| Ketamine vs Thiopental vs Placebo | 20 | **100%** (20) | **50%** (10) | **60%** (12) | **65%** (13) |
| Ketamine vs Placebo | 20 | **95%** (19) | **60%** (12) | 5% (1) | **65%** (13) |
| Thiopental vs Placebo | 20 | **65%** (13) | 30% (6) | **90%** (18) | **75%** (15) |
| Ketamine vs Thiopental | 20 | **100%** (20) | 0% (0) | 5% (1) | **50%** (10) |
| All Comparisons | 80 | **90%** (72) | 35% (28) | 40% (32) | **63.75%** (51) |

The table presents the EEG biomarker categories that were selected in at least half ( $\geq 10/20$ ) of the cross-validation folds for at least one of the drug condition comparisons listed in the first column. For each comparison, the values to the right indicate the percentage of folds in which each category was selected, with the number of selections in parentheses (20 folds total: 5 per alpha level). Bolded numbers identify the categories selected in at least 50% of the folds for each comparison. The final row shows the number of selections (80) across all the comparisons.

##### Relationship between selected EEG biomarkers and behavior

The feature selection algorithm identified four EEG biomarker categories, comprising 24 individual biomarkers. Of these, 13 biomarkers belonging to the PSD exponent, Beta power (total), and Peak delta Power (periodic) categories were significantly (corrected) associated with the CADSS total score (log_10_-transformed). The strongest associations were observed for members of the PSD exponent category (largest β=-0.550, *p*_Adj_<0.0001), followed by members of the beta power (total) category (largest β=-0.454, *p*_Adj_<0.0001), and a member of the peak delta power (periodic) category (largest β=-0.308, *p*_Adj_=0.0022). For the HVLT total score, four biomarkers belonging to the PSD exponent category showed significant (corrected) associations (largest β=-0.309, *p*_Adj_< 0.0202). For the full list of the regression results, see Supplementary Table S11.

## Discussion

In this randomized, placebo-controlled study, we examined the effects that transient, pharmacologically induced shifts in cortical E/I balance have on the aperiodic component of EEG signals, and how these effects relate to subjective drug experiences and cognitive performance. We assessed the aperiodic component by estimating the exponent of the power spectral density (PSD) from EEG signals collected during an oddball task. To manipulate E/I balance, we intravenously administered subanesthetic doses of ketamine and thiopental, agents with comparable sedative profiles known to transiently increase and decrease E/I ratio, respectively. In addition, we used a data-driven, information theory-based approach to evaluate how information relevant for differentiating between drug conditions was carried by the PSD exponent compared to a broad range of commonly used EEG and MEG measures.

We found that under ketamine, the PSD exponent was reduced (i.e., flatter PSD curve) compared to both placebo and thiopental, and that under thiopental, it was increased relative to placebo (i.e., steeper PSD curve). These effects were consistent across absolute and relative (change from pre-infusion) values, and across the pre-stimulation and stimulation periods of the oddball task. As expected, both ketamine and thiopental increased perceptual alterations compared to placebo, with ketamine increasing them more than thiopental. The regression analyses revealed that in the active drug conditions, lower PSD exponent values were associated with greater subjective effects. Similarly, ketamine and thiopental induced verbal memory deficits compared to placebo, with ketamine inducing larger deficits than thiopental. Regression analyses also showed that lower PSD exponent values were associated with greater disruptions in verbal memory performance.

Our data-driven analyses identified four EEG biomarker categories that were consistently selected in at least one drug condition comparison. These results suggest that among all the biomarker categories, these ones carried the most relevant information for distinguishing between drug conditions under selection size constraints. Among these, the PSD exponent was the most consistently selected category across comparisons, suggesting that it carried a comparatively greater amount of useful information for distinguishing drug conditions. Total beta power ranked second in overall consistency and therefore in the amount of useful information it carried. In contrast, in the pairwise comparisons, peak delta power was informative for differentiating between ketamine and placebo only, while total theta power was informative for differentiating between thiopental and placebo only.

Our results in awake humans showed that a transient, drug-induced increase in E/I balance reduced the PSD exponent, whereas a transient, drug-induced reduction in E/I balance increased it. These findings provide the first evidence in awake humans, using drugs with comparable sedative effects, to support recent modeling work proposing an inverse relationship between E/I balance in cortical circuits and the PSD exponent of global electrophysiological signals^29^. While these results suggest that the PSD exponent may serve as a noninvasive measure sensitive to shifts in E/I balance in humans, especially in within-subject designs, they must be replicated using other compounds with known effects on E/I balance to assess their generalizability and limitations.

A wide range of EEG biomarkers commonly used to characterize drug-induced changes in brain dynamics failed to reach the consistency (informativeness) threshold we set in our feature selection algorithm. As a result, well-established measures such as signal-to-noise ratio and Lempel-Ziv complexity were excluded by our approach. Importantly, these results do not imply that biomarker categories that were not selected by our algorithm captured no relevant or unique drug-related information, but that under pressure to differentiate between the drug conditions with as few biomarkers as possible, the selected ones stood out as carrying the largest amount of relevant information.

Variability in the selected biomarkers across pairwise comparisons suggests that some variables carried information about drug-induced changes in E/I balance that was not fully captured by the changes in the PSD exponent. Specifically, total beta power was selected as informative in every pairwise comparison of drug conditions. While the interpretation of these findings goes beyond the scope of this article, which focuses on the interaction between E/I balance and the aperiodic component, it is important to highlight that total beta power was considered informative across comparisons. This coincides with the well-known effects of thiopental increasing^45^ and ketamine suppressing^46, 47^ power in the beta band. In addition, periodic delta power was useful for distinguishing ketamine from placebo, and total theta power was useful for distinguishing thiopental from placebo. The value of these measures as informative biomarkers to differentiate between states of increased and decreased E/I balance is something that will need to be explored in future studies.

Ketamine, an NMDA receptor antagonist, produces dissociative anesthesia at higher doses (2mg/kg) and rapid antidepressant effects at lower doses (0.5-1.0mg/kg)^48, 49^. In this study, a subanesthetic reduced the PSD exponent, supporting the hypothesis that ketamine’s disinhibitory effects are associated with a shift in EEG spectral power from lower to higher frequencies^50^. Preclinical studies have shown that ketamine increases E/I balance in the medial prefrontal cortex (mPFC) by reducing interneuron activity, as demonstrated using electrophysiology^51–53^, microdialysis^54^, and other approaches^55^. However, intracortical recordings in nonhuman primates have shown that ketamine’s effects vary across neuronal populations and cortical layers^56^. While ketamine disinhibits pyramidal cells in cortical layer 5 (L5), it inhibits them in cortical layer 3 (L3)^56^. Interestingly, recent modeling work using biophysically realistic neurons showed that L5 pyramidal cells are by far the largest contributor to EEG signals^57^. Therefore, our results showing a reduction in the PSD exponent, which is consistent increased E/I balance, is likely reflecting the ketamine’s disinhibitory effects on L5, as its inhibitory effects on L3 are likely masked^56^.

Thiopental acts by enhancing GABA-mediated inhibition of synaptic transmission, leading to a decrease in E/I balance. In this study, we observed an increase in the PSD exponent following thiopental administration, consistent with preclinical and human studies showing that GABAergic agents increase the PSD exponent^27, 30^.

At subanesthetic doses, ketamine and related NMDA receptor antagonists increase the E/I ratio by disinhibiting cortical circuits, while thiopental and other GABAergic agents reduce the E/I ratio by enhancing inhibition^20, 39, 51–53^. Despite having opposite effects on E/I balance, they both induce significant impairment in brain function^58–61 62^, suggesting an inverted-U relationship between E/I balance and brain circuit output. Deviations in either direction from an optimal task-related E/I balance would lead to reduced neural efficiency^3–6^. The nature of the impairment depends on the direction of the shift in E/I balance. Computational models suggest that elevated E/I balance can lead to impulsive responses, while reduced E/I balance leads to indecisive responses in decision-making tasks^6^. Consistently, in this study, we observed that both ketamine-induced reductions (increased E/I ratio) and thiopental-induced increases (reduced E/I ratio) in the PSD exponent were associated with increased subjective effects and verbal memory impairment. This is consistent with previous studies showing that the PSD exponent tracks neurocognitive performance across domains^27, 63, 64^, including verbal learning tasks^65^. However, our results also showed that ketamine’s subjective and cognitive effects were stronger than those of thiopental, indicating an asymmetry in the inverted-U relationship. Thus, at comparable sedation levels, the substances showed that circuit disinhibition induced more severe subjective and cognitive disruptions than hyperinhibition.

The subjective effects of ketamine have been linked to its antidepressant action, such that higher CADSS scores at 40 minutes post-infusion correlate with reductions in depressive symptoms at both 230 minutes and 7 days post-infusion. Considering the positive correlation between CADSS scores and PSD exponent observed in this study, the association between CADSS scores and antidepressant effects raises the possibility that the PSD exponent may help identify patients who are more likely to benefit from ketamine treatment. If future studies confirm this relationship, it could eventually help reduce the economic, social, and personal burden associated with current trial-and-error treatment strategies.

There are several limitations to this study. It is important to note that E/I balance can be conceptualized at various scales, from excitatory and inhibitory inputs onto a single neuron to long-range neural networks^66^. The global scale of EEG signals cannot disentangle the specific changes occurring at the cellular or neuroreceptor level. Other approaches, including magnetic resonance spectroscopy (MRS)^67^ and neuroimaging^10, 68^, as well as transcranial magnetic stimulation (TMS) paradigms, such as short-interval intracortical inhibition (SICI), intracortical facilitation (ICF), and long-interval intracortical inhibition (LICI)^69^, offer alternative markers of E/I balance (e.g., receptor occupancy, neurotransmitter metabolism, and concentration) in humans. For example, pharmacological fMRI has been used to generate empirical models of E/I balance based on functional connectivity^70, 71^. Furthermore, cerebrospinal fluid GABA levels have been used as a proxy measure for E/I balance alterations in patients with schizophrenia^72–74^. However, these methods are relatively intrusive, expensive, and/or require specialized infrastructure. Additionally, their poor temporal resolution makes it difficult to capture the fast dynamics of E/I balance. Nevertheless, parallel studies using these approaches in humans in combination with pharmacological agents with known effects on E/I balance may help to better understand and identify the electrophysiological fingerprints of illness- or drug-induced alterations in E/I balance. Currently, there is a range of promising electrophysiological measures such as microstates, entropy, and beta and gamma power^1, 75^.

While the changes in the PSD exponent induced by ketamine and thiopental were consistent with their effects on E/I balance^51–53^, we cannot conclusively rule out the contribution of factors unrelated to excitation and inhibition, such as drug-induced changes in the biophysical properties of neurons^76^. Furthermore, ketamine affects multiple neurotransmitter systems, including the opioid, aminergic, and cholinergic systems^77^, making it difficult to determine how each pathway contributes to changes in the electrophysiological signals.

A further limitation concerns our data-driven biomarker selection approach, which may disregard mechanistically relevant measures. The method is designed to maximize mutual information with the drug condition labels while minimizing the size of the selected biomarker set. As a result, when multiple biomarkers carry overlapping information, the algorithm tends to retain only the most informative one, excluding others that may be less informative but still relevant for understanding the neural mechanisms underlying the effects of ketamine and thiopental.

## Methods

### Participants

91 healthy adults (44 female; age = 24.220 ± 2.590 years) participated in this double-blind, placebo-controlled, within-subject, three-test-day challenge study. Participants were recruited through public advertisement and provided written informed consent prior to participation. All participants were medically and neurologically healthy and underwent psychiatric screening using the Structured Clinical Interview for DSM-IV. Individuals with a history of alcohol abuse or dependence, other Axis I diagnoses, or a family history of psychosis were excluded. Participants remained alcohol-free for three days before each test day and throughout the study, and completed urine toxicology and breathalyzer tests on each test day. To reduce the risk of nausea, participants were asked to fast the night before each test day and during each session. The study was approved by the Human Subjects Subcommittee of the VA Connecticut Healthcare System (West Haven, CT, USA) and the Human Investigations Committee of Yale University School of Medicine (New Haven, CT, USA). Participant safety was reviewed on an ongoing basis by the Data Safety Monitoring Board of the NIAAA Center for the Translational Neuroscience of Alcoholism.

### Drug challenge sessions

Each participant received a 60-min intravenous (IV) infusion of saline (placebo) and ketamine (0.23 mg/kg loading and 0.58 mg/kg/hr) administered on separate test days in randomized order. A subset of 63 participants also received a 60-minute IV infusion of thiopental (1.5mg/kg loading and 40 mcg/kg/minute). Consecutive sessions were separated by ≥ 3 days. Doses were selected to have mild-to-moderate psychogenic and sedating effects based on previous studies ^20, 78, 79^.

### Behavioral measures

Psychoactive effects were assessed using the Clinician Administered Dissociative States Scale (CADSS)^80^, and verbal memory performance was assessed using the Hopkins Verbal Learning Test (HVLT)^81^. These measures were selected because they have been shown to be sensitive to the effects of low doses of ketamine^82, 83^.

### EEG paradigm

EEG data were collected while participants sat in a dimly lit, sound-attenuated booth and completed two runs of a three-stimulus visual oddball task. The first run occurred approximately 90 minutes before infusion, and the second run 45 minutes after the start of infusion. Each run comprised three blocks of 150 stimuli presented in pseudo-random order on a computer screen. Stimuli included standard (small blue circle, white background; 80%), target (large blue circle, white background; 10%), and novel (non-repeating fractal images; 10%) images, each presented for 500 ms with a 1500 ms inter-stimulus interval. Participants were instructed to respond only to target stimuli by pressing a button with their preferred hand.

### EEG acquisition

EEG data were acquired at a sampling rate of 1000Hz using sintered electrodes and a Compumedics Neuroscan EEG system. Recordings were obtained from three midline scalp sites (Fz, Cz, and Pz) using a linked-earlobe reference and a fronto-central ground. Vertical and horizontal electrooculograms (EOGs) were recorded using electrodes placed above and below the right eye, and on the outer canthi of both eyes, respectively. Electrode impedances did not exceed 10 kΩ.

### EEG data processing and analysis

#### EEG preprocessing

Continuous EEG and EOG signals were bandpass (0.5-100Hz) and notch filtered (59-61Hz) using a windowed sinc FIR filter (−6dB/Hz). Eye movement artifacts were corrected using a standard regression procedure^84^. Data were segmented in 1250ms epochs time-locked to stimulus onset, with a 400ms pre-stimulus baseline. Trials containing voltages exceeding ±100 *μ*V or muscular artifacts were excluded from analysis.

#### EEG analysis

EEG biomarkers were extracted separately for each subject, electrode, and drug condition. To better capture the effects of drugs, the EEG biomarkers were calculated in both absolute and relative terms, i.e., for the post-infusion period and as a change from pre-infusion, respectively. Finally, the outcome variables were averaged across electrodes Cz and Pz. Fz was left out of the analyses, as Cz and Pz are the least affected by motor artifacts^85^.

##### Aperiodic component estimation

Due to the structure of the oddball task, 80% of trials consisted of standard stimuli. To increase the reliability of the PSD exponent (*χ*), we computed the PSD using only standard trials, allowing trial-wise PSD fluctuations to be attenuated through averaging. For each subject, electrode, and drug condition, PSDs were computed for each trial’s pre-stimulus baseline (− 400ms to –1ms) and post-stimulus (1ms to 850ms) periods using Welch’s method. The resulting pre- and post-stimulus PSDs were averaged across trials and fitted with the Python implementation of the Fitting Oscillations & One Over F (FOOOF) algorithm^86^ (https://fooof-tools.github.io/fooof/) within the 20-50Hz range, where the correlation between E/I ratio and PSD exponent is strongest^29^. This algorithm has been shown to be robust and has been validated in both animals and humans^87, 88^. The algorithm models the log-log power spectrum as the sum of a linear aperiodic component and multiple Gaussian peaks that capture periodic, oscillatory activity. The aperiodic component is defined by an offset and a spectral exponent *χ*, which describes the steepness of the PSD’s decay with frequency. Oscillatory peaks are modelled as Gaussian curves above the aperiodic fit with peak power, center frequency, and bandwidth as parameters. The algorithm was run with its default parameters and constrained to fit no more than two peaks. The aperiodic PSD exponent χ was used in the hypothesis-driven analyses, while features of the periodic component (peak power, center frequency, and bandwidth) were used in the data-driven portion of the study.

##### Conventional EEG biomarkers

For a complete list of the EEG biomarkers used for comparison, see Supplementary Table S1. Briefly, the cleaned EEG signals were bandpass filtered using windowed sinc FIR filters (− 6dB/Hz) and separated into the following frequency bands: delta (0.5-4Hz), theta (4-8Hz), alpha (8-12Hz), beta (12-30Hz), gamma (30-55Hz), gamma 2 (65-100Hz), and full spectrum (0.5-100Hz). The gamma band was divided into two segments to avoid the distorting effects around 60 Hz caused by the notch filter used to suppress AC line noise.

Based on the inspection of the grand-average of the full spectrum signal, the bandpass-filtered data were segmented into the following task-relevant time windows: pre-stimulus baseline (−250 to −1ms), post-stimulus N100 window (1-250ms), P300 windows (251-500ms), and full stimulation interval (1-500ms). The baseline window was set to 250 ms to match the length of the ERP analysis windows and facilitate comparison. For each drug condition, all biomarkers were computed as both absolute (post-infusion) and relative (change from pre-infusion) values.

###### Event-related potentials (ERPs)

For each electrode, full-spectrum signals were averaged across trials on a point-by-point basis to obtain the ERP waveform. The resulting waveform was baseline-corrected by subtracting the mean voltage of the pre-stimulus baseline period. The N100 and P300 peak amplitudes were defined as the most negative voltage within the 1-250ms post-stimulus interval and the most positive voltage within the 251-500 ms post-stimulus interval, respectively. An automated algorithm was used to detect and extract peak amplitudes and latencies.

###### Total spectral power per frequency band

For each electrode, frequency band (including full spectrum), and time window, total spectral power was estimated by computing the mean squared amplitude of the bandpass filtered signal for each trial and then averaging across trials.

###### Evoked power

For both the N100 and P300 ERPs, evoked power was calculated as the mean squared amplitude of the ERP waveform within the corresponding time window. This value was baseline-corrected by subtracting the mean squared amplitude of the pre-stimulus baseline.

###### Signal-to-noise ratio (SNR)

In this context, the signal refers to the portion of EEG activity that is consistently evoked across trials following stimulus onset, while noise refers to the remaining components of the EEG activity that are not part of the stimulus-locked response. By averaging epochs across trials, non-phase-locked activity cancels out, leaving an approximation of the underlying evoked response, which is reflected in the ERP waveform. Thus, the mean squared amplitude of the ERP within each time window was taken as the signal power. Total power was estimated by calculating the mean squared amplitude for each individual trial and then averaging across trials. Noise power was defined as the difference between total power and signal power. SNR was then computed as the ratio of signal power to noise power.

###### Lempel-Ziv complexity (LZC)

LZC is a non-linear measure originally developed to quantify the randomness of signals^89^. It estimates the minimum number of distinct terms (e.g., “words” or “phonemes”) required to reconstruct a signal (e.g., “sentence”) without information loss. When applied to EEG, LZC reflects the number of distinct activity patterns needed to fully characterize a signal. As the randomness of a signal increases, the number of distinct patterns required also increases, resulting in higher LZC values. For infinitely long white noise-like signals, which lack regular recurrent patterns, the normalized LZC approaches 1, while for infinite regular (periodic) signals it approaches 0 ^90, 91^.

For each electrode and task window, full-spectrum signals were coded into binary sequences using a standard thresholding approach^92^. Values above the median were assigned a 1, and values at or below were assigned a 0^93^. This binarization method is robust to outliers and preserves the relative frequency structure of the signal^92^. For each trial and task window, the binary sequence was parsed using Lempel’s and Ziv’s exhaustive algorithm to identify the minimal set of non-redundant symbolic substrings needed to reconstruct the sequece^89^. The raw complexity count was then normalized by dividing by *n*⁄log_2_ *n*, the theoretical upper bound of LZC as the sequence length *n* approaches infinity. For each electrode and task window, LZC values were averaged across trials.

### Identification of informative EEG biomarkers for differentiating drug conditions

#### Mutual information-based approach to identify EEG biomarkers

We used a custom feature selection algorithm based on mutual information (MI) to identify the subsets of EEG biomarkers that carried the most information about the drug conditions^94^. The algorithm jointly maximizes the MI between sets of selected biomarkers and the drug condition labels while minimizing set sizes. Thus, this approach identifies compact sets of biomarkers that are maximally informative under set size constraints.

The optimization of the algorithm balances two competing objectives that, typically, pull in opposite directions: maximizing information content and minimizing subset size. A regularization parameter, alpha, governs this tradeoff by penalizing larger sets. Alpha can be interpreted as a selection set size constraint, with higher alpha values resulting in more compact subsets. To examine the stability and informativeness of individual biomarkers and biomarker categories across subset sizes, the algorithm was run independently at four levels of alpha: 0.01 (weak constraint), 0.1 (moderate), 0.2 (strong), and 0.5 (very strong).

#### EEG biomarker identification and testing

The dataset was partitioned using a 5-fold cross-validation scheme. Subjects were partitioned into five mutually exclusive groups balanced in size. Each group was used once as the test set, while the remaining four served as the training set, resulting in five independent train-test splits. For both the 3- and 2-drug condition comparisons, the algorithm was trained independently under four levels of the selection size restriction parameter alpha: 0.01 (weak constraint), 0.1 (moderate), 0.2 (strong), and 0.5 (very strong).

The informativeness of the selected EEG biomarker sets was evaluated on each test set independently using random forest classifiers with 100 trees, trained on the remaining 4 groups not used for the evaluation (training set). Classifier performance was reported as the average and standard deviation across testing sets of accuracy, precision, and sensitivity (recall) in the drug condition classification task. Accuracy refers to the percentage of correctly classified cases out of the total number of cases; precision refers to the percentage of correct classifications for a given drug condition out of all the cases predicted as that drug; and sensitivity refers to the percentage of correct classifications for a given drug condition out of all the cases that actually received that drug. The number of EEG biomarkers selected at each alpha level was reported as the average across training folds.

#### EEG biomarker categories

Closely related EEG biomarkers (e.g., PSD exponent computed from pre- and post-stimulus onset windows) contain overlapping information. This overlap may cause the feature selection algorithm to retain only the most informative member of each biomarker family, discarding related biomarkers as redundant. In consequence, informative biomarkers could remain unselected, masked by measures with partially redundant information. To address this issue, we grouped related biomarkers into categories, and report biomarker categories rather than individual biomarkers. For a complete list of the EEG biomarkers within each category, see Supplementary Table S1. For each drug condition comparison, irrespective of how many individual biomarkers from a category were selected in any given fold, each category was counted only once in that fold. This approach ensures clarity about the type of biomarker information that is relevant for differentiating between drug conditions. To reduce the chances of EEG biomarker categories being selected by chance due to inhomogeneous data distributions across the cross-validation splits, the final set of EEG biomarker categories used for the correlation analyses with behavior (see below) was restricted to categories that were consistently informative across the five cross-validation folds and four alpha levels. Specifically, for each drug comparison, there were 20 selection opportunities (5 folds × 4 alpha levels). EEG biomarkers used for the correlation analyses were those belonging to biomarker categories selected in at least 50% (≥10 out of 20) of these selection opportunities.

### Statistical analyses

The data were first examined descriptively using means, standard deviations, histograms, and box plots. Outcome measures were tested for normality using the Kolmogorov-Smirnov test. To evaluate the effect of drug condition on the absolute (post-infusion) and relative (change from pre-infusion) PSD exponent values computed for the pre- and post-stimulus onset windows, Generalized Estimating Equations (GEE) models were used, specifying an unstructured working correlation matrix and drug condition (placebo, thiopental, ketamine) as a within-subject factor. GEE modeling is a robust statistical approach that accounts for within-subject correlations and handles missing data^95^. Significant main effects were followed by post hoc pairwise comparisons, with *p*-values adjusted using the Holm-Bonferroni procedure. Due to floor effects and positive skewness of the CADSS scores in the placebo condition and ceiling effects for the HVLT average scores, behavioral measures were analyzed using a nonparametric approach for longitudinal data^96^. Specifically, the data were ranked and fitted with a mixed-effects model using an unstructured variance-covariance matrix and drug condition as within-subject factor; *p*-values were adjusted for analysis of variance-type statistics. In case of a significant main effect of drug condition, pairwise comparisons were conducted, and *p*-values were adjusted with the Holm-Bonferroni procedure.

Longitudinal GEE regression models with an unstructured working correlation matrix were used to examine the relationships between the PSD exponent and both the subjective (CADSS) and cognitive (HVLT) effects during the active drug conditions. Specifically, the CADSS total (log_10_-transformed) and HVLT average scores were regressed separately on the PSD exponent, estimated as both absolute (post-infusion) and relative (change from pre-infusion) values for the pre-stimulation and stimulation periods of the oddball task. Analyses were conducted on data pooled across the ketamine and thiopental conditions. Standardized regression coefficients (β) were obtained by standardizing the variables prior to fitting the models. For the CADSS and HVLT scores, the p-values were corrected using the Holm-Bonferroni procedure for 4 comparisons each. Marginal R^2^ values were calculated to assess the proportion of variance explained by each GEE model, following the method proposed by Zheng ^97^. Unless otherwise specified, all statistical analyses were conducted using SPSS 29 (IBM Corporation, Armonk, New York) and the nparLD package for R^98^ in RStudio 2024.12.1+563^99^.

#### Relationship between selected EEG biomarker categories and behavioral outcomes

The same longitudinal GEE regression models used to explore the relationship between the PSD exponent and behavioral outcomes were used to examine the relationship between each selected EEG biomarker category and both the subjective (CADSS) and cognitive (HVLT) effects of the active drug conditions. Standardized regression coefficients (β) and marginal R^2^ values were obtained. P-values were corrected using the Holm-Bonferroni procedure for the total number of EEG biomarkers included across all selected categories, separately for the CADSS and HVLT scores.

## Acknowledgements

The authors acknowledge the important contributions of Angelina Genovese, R.N.C., M.B.A., Elizabeth O’Donnell, R.N., and Brenda Breault, R.N., B.S.N., of the Neurobiological Studies Unit of the VA Connecticut Healthcare System, West Haven, CT. In addition, the authors acknowledge support for this work from the Department of Veterans Affairs (VA Merit Review Grant, Alcohol Research Center), National Institute on Alcohol Abuse and Alcoholism (KO5 AA 14906-04, 2P50-AA012870-07, T-32 AA 015496-02), and ANID-Chile Millennium Science Initiative Program (ICN2021-004 to P.A.E.). Finally, this manuscript is dedicated to the memory of our dear friend and colleague the late Dr. Patrick D. Skosnik.

