## Supplementary Tables S2 to S10 for "Opposing Modulation of EEG Aperiodic Component by Ketamine and Thiopental: Implications for the Noninvasive Assessment of Cortical E/I Balance in Humans"

### Supplementary Tables S2-S5: selected EEG biomarkers

#### S2. Supplementary Tables: selected EEG biomarkers for each drug comparison (all alpha values combined)

##### Supplementary Table S2.1. selected EEG biomarkers for the ketamine, thiopental, and placebo comparison (all alpha values combined)

| EEG biomarker | # of selections | % of selections |
|---|---|---|
| avg_standards_20to50hz_post_stim_post_sub_pre_slope | 20 | 100.00% |
| Pow_1-250ms_betaHz_PostSubPre | 13 | 65.00% |
| Pow_1-500ms_thetaHz_PostSubPre | 10 | 50.00% |
| avg_standards_delta_pre_stim_post_sub_pre_peak_pw_0s | 10 | 50.00% |
| EvokedPow_251_500ms_Full_spect_PostInf | 9 | 45.00% |
| Pow_1-500ms_betaHz_PostSubPre | 5 | 25.00% |
| Pow_-250--1ms_betaHz_PostSubPre | 4 | 20.00% |
| Pow_1-250ms_thetaHz_PostSubPre | 2 | 10.00% |
| avg_standards_20to50hz_pre_stim_post_sub_pre_slope | 2 | 10.00% |
| Pow_1-500ms_alphaHz_PostSubPre | 2 | 10.00% |
| avg_standards_20to50hz_post_stim_post_inf_slope | 2 | 10.00% |
| Pow_1-250ms_betaHz_PostInf | 1 | 5.00% |
| Peak_raw_251_500ms_Full_spect_PostSubPre | 1 | 5.00% |
| SNR_rms_raw_251_500ms_Full_spect_PostInf | 1 | 5.00% |
| avg_standards_beta_pre_stim_post_sub_pre_peak_cf_0s | 1 | 5.00% |
| Pow_-250--1ms_gamma2Hz_PostSubPre | 1 | 5.00% |
| Peak_raw_251_500ms_Full_spect_PostInf | 1 | 5.00% |
| Pow_251-500ms_betaHz_PostSubPre | 1 | 5.00% |
| avg_standards_alpha_post_stim_post_sub_pre_peak_cf_0s | 1 | 5.00% |
| Pow_-250--1ms_betaHz_PostInf | 1 | 5.00% |
| Pow_-250--1ms_thetaHz_PostSubPre | 1 | 5.00% |
| avg_standards_gammato50_post_stim_post_sub_pre_peak_pw_0s | 1 | 5.00% |
| avg_standards_alpha_post_stim_post_sub_pre_peak_pw_1s | 1 | 5.00% |
| LZC_Sig_251-500ms_fullspectHz_PostSubPre | 1 | 5.00% |
| avg_standards_theta_pre_stim_post_sub_pre_peak_pw_0s | 1 | 5.00% |
| avg_standards_beta_post_stim_post_sub_pre_peak_pw_1s | 1 | 5.00% |
| Pow_251-500ms_deltaHz_PostInf | 1 | 5.00% |

**Supplementary Table S2.2. selected EEG biomarkers for the ketamine and placebo comparison (all alpha values combined)**

| EEG biomarker | # of selections | % of selections |
|---|---|---|
| avg_standards_20to50hz_post_stim_post_sub_pre_slope | 19 | 95.00% |
| avg_standards_delta_pre_stim_post_sub_pre_peak_pw_0s | 11 | 55.00% |
| Pow_1-250ms_betaHz_PostSubPre | 9 | 45.00% |
| avg_standards_20to50hz_post_stim_post_inf_slope | 6 | 30.00% |
| Pow_251-500ms_deltaHz_PostInf | 6 | 30.00% |
| Pow_-250--1ms_betaHz_PostSubPre | 6 | 30.00% |
| Pow_1-500ms_betaHz_PostSubPre | 6 | 30.00% |
| Peak_raw_251_500ms_Full_spect_PostSubPre | 5 | 25.00% |
| avg_standards_delta_post_stim_post_inf_peak_pw_1s | 4 | 20.00% |
| Pow_1-500ms_alphaHz_PostSubPre | 3 | 15.00% |
| avg_standards_alpha_post_stim_post_inf_peak_cf_0s | 3 | 15.00% |
| avg_standards_beta_pre_stim_post_sub_pre_peak_bw_0s | 3 | 15.00% |
| avg_standards_beta_post_stim_post_sub_pre_peak_pw_1s | 3 | 15.00% |
| avg_standards_beta_post_stim_post_inf_peak_bw_1s | 3 | 15.00% |
| Pow_1-500ms_deltaHz_PostInf | 2 | 10.00% |
| avg_standards_alpha_post_stim_post_sub_pre_peak_cf_0s | 2 | 10.00% |
| avg_standards_beta_pre_stim_post_inf_peak_cf_0s | 2 | 10.00% |
| avg_standards_delta_pre_stim_post_inf_peak_pw_0s | 2 | 10.00% |
| avg_standards_delta_post_stim_post_sub_pre_peak_cf_1s | 2 | 10.00% |
| avg_standards_alpha_post_stim_post_sub_pre_peak_pw_1s | 2 | 10.00% |
| avg_standards_20to50hz_pre_stim_post_inf_slope | 2 | 10.00% |
| Peak_raw_251_500ms_Full_spect_PostInf | 2 | 10.00% |
| avg_standards_gammato50_post_stim_post_inf_peak_pw_0s | 1 | 5.00% |
| avg_standards_delta_post_stim_post_sub_pre_peak_pw_0s | 1 | 5.00% |
| EvokedPow_251_500ms_Full_spect_PostSubPre | 1 | 5.00% |
| Pow_1-250ms_thetaHz_PostSubPre | 1 | 5.00% |
| avg_standards_delta_pre_stim_post_sub_pre_peak_cf_0s | 1 | 5.00% |
| EvokedPow_251_500ms_Full_spect_PostInf | 1 | 5.00% |
| Pow_251-500ms_deltaHz_PostSubPre | 1 | 5.00% |
| Pow_-250--1ms_alphaHz_PostSubPre | 1 | 5.00% |
| Pow_-250--1ms_deltaHz_PostInf | 1 | 5.00% |
| avg_standards_gammato50_pre_stim_post_inf_peak_pw_1s | 1 | 5.00% |
| avg_standards_beta_post_stim_post_sub_pre_peak_bw_0s | 1 | 5.00% |
| avg_standards_20to50hz_pre_stim_post_sub_pre_slope | 1 | 5.00% |
| avg_standards_gammato50_pre_stim_post_sub_pre_peak_pw_0s | 1 | 5.00% |
| avg_standards_theta_pre_stim_post_sub_pre_peak_bw_0s | 1 | 5.00% |
| avg_standards_gammato50_post_stim_post_sub_pre_peak_cf_1s | 1 | 5.00% |
| avg_standards_beta_pre_stim_post_sub_pre_peak_bw_1s | 1 | 5.00% |
| avg_standards_alpha_post_stim_post_sub_pre_peak_cf_1s | 1 | 5.00% |
| Pow_1-250ms_alphaHz_PostSubPre | 1 | 5.00% |

**Supplementary Table S2.3. selected EEG biomarkers for the thiopental and placebo comparison (all alpha values combined)**

| EEG biomarker | # of selections | % of selections |
|---|---|---|
| Pow_1-500ms_thetaHz_PostSubPre | 13 | 65.00% |
| avg_standards_20to50hz_post_stim_post_sub_pre_slope | 12 | 60.00% |
| Pow_1-250ms_betaHz_PostSubPre | 11 | 55.00% |
| Pow_-250--1ms_betaHz_PostSubPre | 9 | 45.00% |
| Pow_251-500ms_thetaHz_PostSubPre | 9 | 45.00% |
| LZC_Sig_251-500ms_fullspectHz_PostSubPre | 7 | 35.00% |
| avg_standards_delta_pre_stim_post_sub_pre_peak_pw_0s | 6 | 30.00% |
| avg_standards_20to50hz_pre_stim_post_sub_pre_slope | 6 | 30.00% |
| EvokedPow_251_500ms_Full_spect_PostInf | 6 | 30.00% |
| SNR_rms_raw_251_500ms_Full_spect_PostInf | 5 | 25.00% |
| Pow_251-500ms_thetaHz_PostInf | 4 | 20.00% |
| Pow_1-500ms_betaHz_PostSubPre | 3 | 15.00% |
| LZC_Sig_251-500ms_fullspectHz_PostInf | 3 | 15.00% |
| avg_standards_theta_pre_stim_post_sub_pre_peak_cf_0s | 3 | 15.00% |
| Pow_251-500ms_deltaHz_PostSubPre | 3 | 15.00% |
| Peak_raw_251_500ms_Full_spect_PostInf | 3 | 15.00% |
| SNR_rms_raw_251_500ms_Full_spect_PostSubPre | 2 | 10.00% |
| avg_standards_20to50hz_pre_stim_post_inf_slope | 2 | 10.00% |
| Pow_1-250ms_thetaHz_PostInf | 2 | 10.00% |
| Pow_-250--1ms_alphaHz_PostSubPre | 2 | 10.00% |
| Peak_raw_1_250ms_Full_spect_PostInf | 1 | 5.00% |
| avg_standards_alpha_post_stim_post_sub_pre_peak_pw_0s | 1 | 5.00% |
| Pow_1-250ms_thetaHz_PostSubPre | 1 | 5.00% |
| avg_standards_20to50hz_post_stim_post_inf_slope | 1 | 5.00% |
| avg_standards_delta_pre_stim_post_inf_peak_cf_0s | 1 | 5.00% |
| avg_standards_theta_pre_stim_post_inf_peak_bw_0s | 1 | 5.00% |
| Pow_1-500ms_alphaHz_PostSubPre | 1 | 5.00% |
| Pow_1-250ms_betaHz_PostInf | 1 | 5.00% |
| Peak_raw_251_500ms_Full_spect_PostSubPre | 1 | 5.00% |
| avg_standards_beta_post_stim_post_inf_peak_pw_0s | 1 | 5.00% |
| avg_standards_delta_pre_stim_post_sub_pre_peak_cf_0s | 1 | 5.00% |
| avg_standards_delta_pre_stim_post_inf_peak_pw_0s | 1 | 5.00% |
| avg_standards_beta_post_stim_post_inf_peak_cf_1s | 1 | 5.00% |
| avg_standards_gammato50_pre_stim_post_sub_pre_peak_pw_1s | 1 | 5.00% |

**Supplementary Table S2.4. selected EEG biomarkers for the ketamine and thiopental comparison (all alpha values combined)**

| EEG biomarker | # of selections | % of selections |
|---|---|---|
| avg_standards_20to50hz_post_stim_post_sub_pre_slope | 20 | 100.00% |
| Pow_1-250ms_betaHz_PostSubPre | 10 | 50.00% |
| Pow_1-500ms_betaHz_PostSubPre | 3 | 15.00% |
| Pow_-250--1ms_betaHz_PostSubPre | 1 | 5.00% |
| Pow_-250--1ms_thetaHz_PostSubPre | 1 | 5.00% |
| Pow_251-500ms_betaHz_PostSubPre | 1 | 5.00% |

#### S3. Supplementary Tables: selected EEG biomarkers for each drug comparison across alpha values

**Supplementary Table S3.1. selected EEG biomarkers for the ketamine, thiopental, and placebo comparison at alpha = 0.01**

| EEG biomarker | # of selections | % of selections |
|---|---|---|
| Pow_1-250ms_betaHz_PostSubPre | 5 | 100.00% |
| avg_standards_20to50hz_post_stim_post_sub_pre_slope | 5 | 100.00% |
| Pow_1-500ms_thetaHz_PostSubPre | 5 | 100.00% |
| EvokedPow_251_500ms_Full_spect_PostInf | 4 | 80.00% |
| Pow_1-500ms_betaHz_PostSubPre | 4 | 80.00% |
| avg_standards_delta_pre_stim_post_sub_pre_peak_pw_0s | 4 | 80.00% |
| Pow_-250--1ms_betaHz_PostSubPre | 3 | 60.00% |
| avg_standards_20to50hz_post_stim_post_inf_slope | 2 | 40.00% |
| Pow_1-500ms_alphaHz_PostSubPre | 2 | 40.00% |
| avg_standards_20to50hz_pre_stim_post_sub_pre_slope | 2 | 40.00% |
| Pow_-250--1ms_thetaHz_PostSubPre | 1 | 20.00% |
| Pow_-250--1ms_gamma2Hz_PostSubPre | 1 | 20.00% |
| Peak_raw_251_500ms_Full_spect_PostInf | 1 | 20.00% |
| Pow_251-500ms_betaHz_PostSubPre | 1 | 20.00% |
| Pow_1-250ms_betaHz_PostInf | 1 | 20.00% |
| Pow_-250--1ms_betaHz_PostInf | 1 | 20.00% |
| avg_standards_theta_pre_stim_post_sub_pre_peak_pw_0s | 1 | 20.00% |
| avg_standards_gammato50_post_stim_post_sub_pre_peak_pw_0s | 1 | 20.00% |
| avg_standards_alpha_post_stim_post_sub_pre_peak_pw_1s | 1 | 20.00% |
| LZC_Sig_251-500ms_fullspectHz_PostSubPre | 1 | 20.00% |
| avg_standards_alpha_post_stim_post_sub_pre_peak_cf_0s | 1 | 20.00% |
| avg_standards_beta_post_stim_post_sub_pre_peak_pw_1s | 1 | 20.00% |
| avg_standards_beta_pre_stim_post_sub_pre_peak_cf_0s | 1 | 20.00% |

**Supplementary Table S3.2. selected EEG biomarkers for the Ketamine, Thiopental, and Placebo Comparison at Alpha = 0.1**

| EEG biomarker | # of selections | % of selections |
|---|---|---|
| Pow_1-250ms_betaHz_PostSubPre | 5 | 100.00% |
| avg_standards_20to50hz_post_stim_post_sub_pre_slope | 5 | 100.00% |
| avg_standards_delta_pre_stim_post_sub_pre_peak_pw_0s | 4 | 80.00% |
| EvokedPow_251_500ms_Full_spect_PostInf | 3 | 60.00% |
| Pow_1-500ms_thetaHz_PostSubPre | 3 | 60.00% |
| Pow_1-250ms_thetaHz_PostSubPre | 2 | 40.00% |
| SNR_rms_raw_251_500ms_Full_spect_PostInf | 1 | 20.00% |
| Pow_1-500ms_betaHz_PostSubPre | 1 | 20.00% |
| Pow_-250--1ms_betaHz_PostSubPre | 1 | 20.00% |
| Peak_raw_251_500ms_Full_spect_PostSubPre | 1 | 20.00% |
| Pow_251-500ms_deltaHz_PostInf | 1 | 20.00% |

**Supplementary Table S3.3. selected EEG biomarkers for the ketamine, thiopental, and placebo comparison at alpha = 0.2**

| EEG biomarker | # of selections | % of selections |
|---|---|---|
| avg_standards_20to50hz_post_stim_post_sub_pre_slope | 5 | 100.00% |
| Pow_1-250ms_betaHz_PostSubPre | 3 | 60.00% |
| Pow_1-500ms_thetaHz_PostSubPre | 2 | 40.00% |
| avg_standards_delta_pre_stim_post_sub_pre_peak_pw_0s | 2 | 40.00% |
| EvokedPow_251_500ms_Full_spect_PostInf | 2 | 40.00% |

**Supplementary Table S3.4. selected EEG biomarkers for the ketamine, thiopental, and placebo comparison at alpha = 0.5**

| EEG biomarker | # of selections | % of selections |
|---|---|---|
| avg_standards_20to50hz_post_stim_post_sub_pre_slope | 5 | 100.00% |

**Supplementary Table S3.5. selected EEG biomarkers for the ketamine and placebo comparison at alpha = 0.01**

| EEG biomarker | # of selections | % of selections |
|---|---|---|
| Pow_1-250ms_betaHz_PostSubPre | 5 | 100.00% |
| avg_standards_delta_pre_stim_post_sub_pre_peak_pw_0s | 5 | 100.00% |
| avg_standards_20to50hz_post_stim_post_sub_pre_slope | 5 | 100.00% |
| avg_standards_delta_post_stim_post_inf_peak_pw_1s | 4 | 80.00% |
| avg_standards_20to50hz_post_stim_post_inf_slope | 4 | 80.00% |
| Pow_251-500ms_deltaHz_PostInf | 4 | 80.00% |
| Pow_-250--1ms_betaHz_PostSubPre | 3 | 60.00% |
| Pow_1-500ms_alphaHz_PostSubPre | 3 | 60.00% |
| Pow_1-500ms_betaHz_PostSubPre | 3 | 60.00% |
| avg_standards_delta_post_stim_post_sub_pre_peak_cf_1s | 2 | 40.00% |
| avg_standards_beta_pre_stim_post_inf_peak_cf_0s | 2 | 40.00% |
| avg_standards_beta_post_stim_post_inf_peak_bw_1s | 2 | 40.00% |
| avg_standards_alpha_post_stim_post_sub_pre_peak_cf_0s | 2 | 40.00% |
| avg_standards_beta_post_stim_post_sub_pre_peak_pw_1s | 2 | 40.00% |
| avg_standards_beta_pre_stim_post_sub_pre_peak_bw_0s | 2 | 40.00% |
| Peak_raw_251_500ms_Full_spect_PostInf | 2 | 40.00% |
| avg_standards_alpha_post_stim_post_sub_pre_peak_pw_1s | 2 | 40.00% |
| avg_standards_alpha_post_stim_post_inf_peak_cf_0s | 2 | 40.00% |
| avg_standards_20to50hz_pre_stim_post_inf_slope | 2 | 40.00% |
| avg_standards_gammato50_pre_stim_post_inf_peak_pw_1s | 1 | 20.00% |
| avg_standards_delta_post_stim_post_sub_pre_peak_pw_0s | 1 | 20.00% |
| EvokedPow_251_500ms_Full_spect_PostSubPre | 1 | 20.00% |
| Pow_1-250ms_thetaHz_PostSubPre | 1 | 20.00% |
| avg_standards_delta_pre_stim_post_sub_pre_peak_cf_0s | 1 | 20.00% |
| EvokedPow_251_500ms_Full_spect_PostInf | 1 | 20.00% |
| Pow_251-500ms_deltaHz_PostSubPre | 1 | 20.00% |
| Pow_-250--1ms_alphaHz_PostSubPre | 1 | 20.00% |
| avg_standards_gammato50_post_stim_post_inf_peak_pw_0s | 1 | 20.00% |
| Pow_1-500ms_deltaHz_PostInf | 1 | 20.00% |
| avg_standards_beta_pre_stim_post_sub_pre_peak_bw_1s | 1 | 20.00% |
| Pow_-250--1ms_deltaHz_PostInf | 1 | 20.00% |
| Peak_raw_251_500ms_Full_spect_PostSubPre | 1 | 20.00% |
| avg_standards_beta_post_stim_post_sub_pre_peak_bw_0s | 1 | 20.00% |
| avg_standards_delta_pre_stim_post_inf_peak_pw_0s | 1 | 20.00% |
| avg_standards_20to50hz_pre_stim_post_sub_pre_slope | 1 | 20.00% |
| avg_standards_gammato50_pre_stim_post_sub_pre_peak_pw_0s | 1 | 20.00% |
| avg_standards_theta_pre_stim_post_sub_pre_peak_bw_0s | 1 | 20.00% |
| avg_standards_gammato50_post_stim_post_sub_pre_peak_cf_1s | 1 | 20.00% |
| avg_standards_alpha_post_stim_post_sub_pre_peak_cf_1s | 1 | 20.00% |
| Pow_1-250ms_alphaHz_PostSubPre | 1 | 20.00% |

**Supplementary Table S3.6 selected EEG biomarkers for the ketamine and placebo comparison at alpha = 0.1**

| EEG biomarker | # of selections | % of selections |
|---|---|---|
| avg_standards_20to50hz_post_stim_post_sub_pre_slope | 5 | 100.00% |
| Pow_1-250ms_betaHz_PostSubPre | 3 | 60.00% |
| avg_standards_delta_pre_stim_post_sub_pre_peak_pw_0s | 3 | 60.00% |
| Pow_-250--1ms_betaHz_PostSubPre | 2 | 40.00% |
| Peak_raw_251_500ms_Full_spect_PostSubPre | 2 | 40.00% |
| Pow_251-500ms_deltaHz_PostInf | 2 | 40.00% |
| avg_standards_beta_pre_stim_post_sub_pre_peak_bw_0s | 1 | 20.00% |
| avg_standards_alpha_post_stim_post_inf_peak_cf_0s | 1 | 20.00% |
| Pow_1-500ms_betaHz_PostSubPre | 1 | 20.00% |
| avg_standards_delta_pre_stim_post_inf_peak_pw_0s | 1 | 20.00% |
| avg_standards_beta_post_stim_post_inf_peak_bw_1s | 1 | 20.00% |
| avg_standards_beta_post_stim_post_sub_pre_peak_pw_1s | 1 | 20.00% |
| avg_standards_20to50hz_post_stim_post_inf_slope | 1 | 20.00% |

**Supplementary Table S3.7. selected EEG biomarkers for the ketamine and placebo comparison at alpha = 0.2**

| EEG biomarker | # of selections | % of selections |
|---|---|---|
| avg_standards_20to50hz_post_stim_post_sub_pre_slope | 4 | 80.00% |
| avg_standards_delta_pre_stim_post_sub_pre_peak_pw_0s | 3 | 60.00% |
| Pow_1-500ms_betaHz_PostSubPre | 2 | 40.00% |
| Peak_raw_251_500ms_Full_spect_PostSubPre | 2 | 40.00% |
| Pow_1-250ms_betaHz_PostSubPre | 1 | 20.00% |
| Pow_-250--1ms_betaHz_PostSubPre | 1 | 20.00% |
| Pow_1-500ms_deltaHz_PostInf | 1 | 20.00% |
| avg_standards_20to50hz_post_stim_post_inf_slope | 1 | 20.00% |

**Supplementary Table S3.8. selected EEG biomarkers for the ketamine and placebo comparison at alpha = 0.5**

| EEG biomarker | # of selections | % of selections |
|---|---|---|
| avg_standards_20to50hz_post_stim_post_sub_pre_slope | 5 | 100.00% |

**Supplementary Table S3.9. selected EEG biomarkers for the thiopental and placebo comparison at alpha = 0.01**

| EEG biomarker | # of selections | % of selections |
|---|---|---|
| Pow_-250--1ms_betaHz_PostSubPre | 5 | 100.00% |
| Pow_1-250ms_betaHz_PostSubPre | 5 | 100.00% |
| SNR_rms_raw_251_500ms_Full_spect_PostInf | 5 | 100.00% |
| avg_standards_delta_pre_stim_post_sub_pre_peak_pw_0s | 5 | 100.00% |
| avg_standards_20to50hz_post_stim_post_sub_pre_slope | 5 | 100.00% |
| LZC_Sig_251-500ms_fullspectHz_PostSubPre | 5 | 100.00% |
| Pow_251-500ms_thetaHz_PostSubPre | 5 | 100.00% |
| Pow_1-500ms_thetaHz_PostSubPre | 4 | 80.00% |
| avg_standards_20to50hz_pre_stim_post_sub_pre_slope | 4 | 80.00% |
| Pow_1-500ms_betaHz_PostSubPre | 3 | 60.00% |
| Pow_251-500ms_thetaHz_PostInf | 3 | 60.00% |
| LZC_Sig_251-500ms_fullspectHz_PostInf | 3 | 60.00% |
| Pow_251-500ms_deltaHz_PostSubPre | 2 | 40.00% |
| avg_standards_20to50hz_pre_stim_post_inf_slope | 2 | 40.00% |
| EvokedPow_251_500ms_Full_spect_PostInf | 2 | 40.00% |
| SNR_rms_raw_251_500ms_Full_spect_PostSubPre | 2 | 40.00% |
| Pow_1-250ms_thetaHz_PostInf | 2 | 40.00% |
| avg_standards_theta_pre_stim_post_sub_pre_peak_cf_0s | 2 | 40.00% |
| Pow_-250--1ms_alphaHz_PostSubPre | 2 | 40.00% |
| Pow_1-500ms_alphaHz_PostSubPre | 1 | 20.00% |
| avg_standards_delta_pre_stim_post_inf_peak_pw_0s | 1 | 20.00% |
| avg_standards_delta_pre_stim_post_sub_pre_peak_cf_0s | 1 | 20.00% |
| avg_standards_beta_post_stim_post_inf_peak_pw_0s | 1 | 20.00% |
| Peak_raw_251_500ms_Full_spect_PostSubPre | 1 | 20.00% |
| Pow_1-250ms_betaHz_PostInf | 1 | 20.00% |
| avg_standards_delta_pre_stim_post_inf_peak_cf_0s | 1 | 20.00% |
| avg_standards_theta_pre_stim_post_inf_peak_bw_0s | 1 | 20.00% |
| Peak_raw_251_500ms_Full_spect_PostInf | 1 | 20.00% |
| avg_standards_20to50hz_post_stim_post_inf_slope | 1 | 20.00% |
| Pow_1-250ms_thetaHz_PostSubPre | 1 | 20.00% |
| avg_standards_alpha_post_stim_post_sub_pre_peak_pw_0s | 1 | 20.00% |
| Peak_raw_1_250ms_Full_spect_PostInf | 1 | 20.00% |
| avg_standards_beta_post_stim_post_inf_peak_cf_1s | 1 | 20.00% |

**Supplementary Table S3.10. selected EEG biomarkers for the thiopental and placebo comparison at alpha = 0.1**

| EEG biomarker | # of selections | % of selections |
|---|---|---|
| avg_standards_20to50hz_post_stim_post_sub_pre_slope | 5 | 100.00% |
| EvokedPow_251_500ms_Full_spect_PostInf | 3 | 60.00% |
| Pow_1-250ms_betaHz_PostSubPre | 3 | 60.00% |
| Pow_1-500ms_thetaHz_PostSubPre | 3 | 60.00% |
| Pow_251-500ms_thetaHz_PostSubPre | 2 | 40.00% |
| LZC_Sig_251-500ms_fullspectHz_PostSubPre | 2 | 40.00% |
| Pow_-250--1ms_betaHz_PostSubPre | 1 | 20.00% |
| avg_standards_delta_pre_stim_post_sub_pre_peak_pw_0s | 1 | 20.00% |
| Peak_raw_251_500ms_Full_spect_PostInf | 1 | 20.00% |
| avg_standards_gammato50_pre_stim_post_sub_pre_peak_pw_1s | 1 | 20.00% |
| avg_standards_20to50hz_pre_stim_post_sub_pre_slope | 1 | 20.00% |
| avg_standards_theta_pre_stim_post_sub_pre_peak_cf_0s | 1 | 20.00% |
| Pow_251-500ms_deltaHz_PostSubPre | 1 | 20.00% |

**Supplementary Table S3.11. selected EEG biomarkers for the thiopental and placebo comparison at alpha = 0.2**

| EEG biomarker | # of selections | % of selections |
|---|---|---|
| Pow_1-500ms_thetaHz_PostSubPre | 4 | 80.00% |
| Pow_-250--1ms_betaHz_PostSubPre | 2 | 40.00% |
| Pow_1-250ms_betaHz_PostSubPre | 2 | 40.00% |
| Peak_raw_251_500ms_Full_spect_PostInf | 1 | 20.00% |
| Pow_251-500ms_thetaHz_PostSubPre | 1 | 20.00% |
| Pow_251-500ms_thetaHz_PostInf | 1 | 20.00% |
| EvokedPow_251_500ms_Full_spect_PostInf | 1 | 20.00% |
| avg_standards_20to50hz_post_stim_post_sub_pre_slope | 1 | 20.00% |

**Supplementary Table S3.12. selected EEG biomarkers for the thiopental and placebo comparison at alpha = 0.5**

| EEG biomarker | # of selections | % of selections |
|---|---|---|
| Pow_1-500ms_thetaHz_PostSubPre | 2 | 40.00% |
| Pow_251-500ms_thetaHz_PostSubPre | 1 | 20.00% |
| Pow_-250--1ms_betaHz_PostSubPre | 1 | 20.00% |
| Pow_1-250ms_betaHz_PostSubPre | 1 | 20.00% |
| avg_standards_20to50hz_pre_stim_post_sub_pre_slope | 1 | 20.00% |
| avg_standards_20to50hz_post_stim_post_sub_pre_slope | 1 | 20.00% |

**Supplementary Table S3.13. selected EEG biomarkers for the ketamine and thiopental comparison at alpha = 0.01**

| EEG biomarker | # of selections | % of selections |
|---|---|---|
| Pow_1-250ms_betaHz_PostSubPre | 5 | 100.00% |
| avg_standards_20to50hz_post_stim_post_sub_pre_slope | 5 | 100.00% |
| Pow_1-500ms_betaHz_PostSubPre | 3 | 60.00% |
| Pow_-250--1ms_betaHz_PostSubPre | 1 | 20.00% |
| Pow_-250--1ms_thetaHz_PostSubPre | 1 | 20.00% |
| Pow_251-500ms_betaHz_PostSubPre | 1 | 20.00% |

**Supplementary Table S3.14. selected EEG biomarkers for the ketamine and thiopental comparison at alpha = 0.1**

| EEG biomarker | # of selections | % of selections |
|---|---|---|
| Pow_1-250ms_betaHz_PostSubPre | 5 | 100.00% |
| avg_standards_20to50hz_post_stim_post_sub_pre_slope | 5 | 100.00% |

**Supplementary Table S3.15. selected EEG biomarkers for the ketamine and thiopental comparison at alpha = 0.2**

| EEG biomarker | # of selections | % of selections |
|---|---|---|
| avg_standards_20to50hz_post_stim_post_sub_pre_slope | 5 | 100.00% |

**Supplementary Table S3.16. selected EEG biomarkers for the ketamine and thiopental comparison at alpha = 0.5**

| EEG biomarker | # of selections | % of selections |
|---|---|---|
| avg_standards_20to50hz_post_stim_post_sub_pre_slope | 5 | 100.00% |

**S4. Supplementary Tables: selected EEG biomarkers across alpha values (all drug comparisons combined)**

**Supplementary Table S4.1. selected EEG biomarkers at alpha = 0.01 (all drug comparisons combined)**

| EEG biomarker | # of selections | % of selections |
|---|---|---|
| Pow_1-250ms_betaHz_PostSubPre | 20 | 100.00% |
| avg_standards_20to50hz_post_stim_post_sub_pre_slope | 20 | 100.00% |
| avg_standards_delta_pre_stim_post_sub_pre_peak_pw_0s | 14 | 70.00% |
| Pow_1-500ms_betaHz_PostSubPre | 13 | 65.00% |
| Pow_-250--1ms_betaHz_PostSubPre | 12 | 60.00% |
| Pow_1-500ms_thetaHz_PostSubPre | 9 | 45.00% |
| EvokedPow_251_500ms_Full_spect_PostInf | 7 | 35.00% |
| avg_standards_20to50hz_post_stim_post_inf_slope | 7 | 35.00% |
| avg_standards_20to50hz_pre_stim_post_sub_pre_slope | 7 | 35.00% |
| Pow_1-500ms_alphaHz_PostSubPre | 6 | 30.00% |
| LZC_Sig_251-500ms_fullspectHz_PostSubPre | 6 | 30.00% |
| Pow_251-500ms_thetaHz_PostSubPre | 5 | 25.00% |
| SNR_rms_raw_251_500ms_Full_spect_PostInf | 5 | 25.00% |
| avg_standards_20to50hz_pre_stim_post_inf_slope | 4 | 20.00% |
| Pow_251-500ms_deltaHz_PostInf | 4 | 20.00% |
| avg_standards_delta_post_stim_post_inf_peak_pw_1s | 4 | 20.00% |
| Peak_raw_251_500ms_Full_spect_PostInf | 4 | 20.00% |
| avg_standards_beta_post_stim_post_sub_pre_peak_pw_1s | 3 | 15.00% |
| avg_standards_alpha_post_stim_post_sub_pre_peak_cf_0s | 3 | 15.00% |
| avg_standards_alpha_post_stim_post_sub_pre_peak_pw_1s | 3 | 15.00% |
| Pow_-250--1ms_alphaHz_PostSubPre | 3 | 15.00% |
| Pow_251-500ms_deltaHz_PostSubPre | 3 | 15.00% |
| Pow_251-500ms_thetaHz_PostInf | 3 | 15.00% |
| LZC_Sig_251-500ms_fullspectHz_PostInf | 3 | 15.00% |
| SNR_rms_raw_251_500ms_Full_spect_PostSubPre | 2 | 10.00% |
| Pow_1-250ms_thetaHz_PostInf | 2 | 10.00% |
| avg_standards_theta_pre_stim_post_sub_pre_peak_cf_0s | 2 | 10.00% |
| Pow_1-250ms_thetaHz_PostSubPre | 2 | 10.00% |
| avg_standards_delta_pre_stim_post_sub_pre_peak_cf_0s | 2 | 10.00% |
| Peak_raw_251_500ms_Full_spect_PostSubPre | 2 | 10.00% |
| avg_standards_delta_pre_stim_post_inf_peak_pw_0s | 2 | 10.00% |
| avg_standards_beta_pre_stim_post_inf_peak_cf_0s | 2 | 10.00% |
| avg_standards_beta_post_stim_post_inf_peak_bw_1s | 2 | 10.00% |
| avg_standards_beta_pre_stim_post_sub_pre_peak_bw_0s | 2 | 10.00% |
| avg_standards_alpha_post_stim_post_inf_peak_cf_0s | 2 | 10.00% |
| Pow_251-500ms_betaHz_PostSubPre | 2 | 10.00% |
| Pow_1-250ms_betaHz_PostInf | 2 | 10.00% |
| Pow_-250--1ms_thetaHz_PostSubPre | 2 | 10.00% |
| avg_standards_delta_post_stim_post_sub_pre_peak_cf_1s | 2 | 10.00% |
| Pow_1-250ms_alphaHz_PostSubPre | 1 | 5.00% |
| avg_standards_gammato50_post_stim_post_inf_peak_pw_0s | 1 | 5.00% |
| EvokedPow_251_500ms_Full_spect_PostSubPre | 1 | 5.00% |
| avg_standards_delta_post_stim_post_sub_pre_peak_pw_0s | 1 | 5.00% |
| avg_standards_delta_pre_stim_post_inf_peak_cf_0s | 1 | 5.00% |
| Peak_raw_1_250ms_Full_spect_PostInf | 1 | 5.00% |

| | | |
|---|---|---|
| avg_standards_alpha_post_stim_post_sub_pre_peak_pw_0s | 1 | 5.00% |
| avg_standards_gammato50_pre_stim_post_inf_peak_pw_1s | 1 | 5.00% |
| avg_standards_theta_pre_stim_post_inf_peak_bw_0s | 1 | 5.00% |
| avg_standards_beta_post_stim_post_inf_peak_pw_0s | 1 | 5.00% |
| Pow_1-500ms_deltaHz_PostInf | 1 | 5.00% |
| avg_standards_theta_pre_stim_post_sub_pre_peak_bw_0s | 1 | 5.00% |
| Pow_-250--1ms_deltaHz_PostInf | 1 | 5.00% |
| avg_standards_beta_post_stim_post_sub_pre_peak_bw_0s | 1 | 5.00% |
| avg_standards_gammato50_pre_stim_post_sub_pre_peak_pw_0s | 1 | 5.00% |
| avg_standards_gammato50_post_stim_post_sub_pre_peak_cf_1s | 1 | 5.00% |
| avg_standards_beta_pre_stim_post_sub_pre_peak_bw_1s | 1 | 5.00% |
| avg_standards_alpha_post_stim_post_sub_pre_peak_cf_1s | 1 | 5.00% |
| avg_standards_beta_pre_stim_post_sub_pre_peak_cf_0s | 1 | 5.00% |
| Pow_-250--1ms_gamma2Hz_PostSubPre | 1 | 5.00% |
| Pow_-250--1ms_betaHz_PostInf | 1 | 5.00% |
| avg_standards_gammato50_post_stim_post_sub_pre_peak_pw_0s | 1 | 5.00% |
| avg_standards_theta_pre_stim_post_sub_pre_peak_pw_0s | 1 | 5.00% |
| avg_standards_beta_post_stim_post_inf_peak_cf_1s | 1 | 5.00% |

**Supplementary Table S4.2. selected EEG biomarkers at alpha = 0.1 (all drug comparisons combined)**

| EEG biomarker | # of selections | % of selections |
|---|---|---|
| avg_standards_20to50hz_post_stim_post_sub_pre_slope | 20 | 100.00% |
| Pow_1-250ms_betaHz_PostSubPre | 16 | 80.00% |
| avg_standards_delta_pre_stim_post_sub_pre_peak_pw_0s | 8 | 40.00% |
| EvokedPow_251_500ms_Full_spect_PostInf | 6 | 30.00% |
| Pow_1-500ms_thetaHz_PostSubPre | 6 | 30.00% |
| Pow_-250--1ms_betaHz_PostSubPre | 4 | 20.00% |
| Peak_raw_251_500ms_Full_spect_PostSubPre | 3 | 15.00% |
| Pow_251-500ms_deltaHz_PostInf | 3 | 15.00% |
| Pow_251-500ms_thetaHz_PostSubPre | 2 | 10.00% |
| LZC_Sig_251-500ms_fullspectHz_PostSubPre | 2 | 10.00% |
| Pow_1-250ms_thetaHz_PostSubPre | 2 | 10.00% |
| Pow_1-500ms_betaHz_PostSubPre | 2 | 10.00% |
| avg_standards_20to50hz_post_stim_post_inf_slope | 1 | 5.00% |
| avg_standards_theta_pre_stim_post_sub_pre_peak_cf_0s | 1 | 5.00% |
| avg_standards_20to50hz_pre_stim_post_sub_pre_slope | 1 | 5.00% |
| avg_standards_gammato50_pre_stim_post_sub_pre_peak_pw_1s | 1 | 5.00% |
| Peak_raw_251_500ms_Full_spect_PostInf | 1 | 5.00% |
| SNR_rms_raw_251_500ms_Full_spect_PostInf | 1 | 5.00% |
| avg_standards_beta_post_stim_post_sub_pre_peak_pw_1s | 1 | 5.00% |
| avg_standards_beta_post_stim_post_inf_peak_bw_1s | 1 | 5.00% |
| avg_standards_delta_pre_stim_post_inf_peak_pw_0s | 1 | 5.00% |
| avg_standards_alpha_post_stim_post_inf_peak_cf_0s | 1 | 5.00% |
| avg_standards_beta_pre_stim_post_sub_pre_peak_bw_0s | 1 | 5.00% |
| Pow_251-500ms_deltaHz_PostSubPre | 1 | 5.00% |

**Supplementary Table S4.3. selected EEG biomarkers at alpha = 0.2 (all drug comparisons combined)**

| EEG biomarker | # of selections | % of selections |
|---|---|---|
| avg_standards_20to50hz_post_stim_post_sub_pre_slope | 15 | 75.00% |
| Pow_1-250ms_betaHz_PostSubPre | 6 | 30.00% |
| Pow_1-500ms_thetaHz_PostSubPre | 6 | 30.00% |
| avg_standards_delta_pre_stim_post_sub_pre_peak_pw_0s | 5 | 25.00% |
| EvokedPow_251_500ms_Full_spect_PostInf | 3 | 15.00% |
| Pow_-250--1ms_betaHz_PostSubPre | 3 | 15.00% |
| Pow_1-500ms_betaHz_PostSubPre | 2 | 10.00% |
| Peak_raw_251_500ms_Full_spect_PostSubPre | 2 | 10.00% |
| Pow_1-500ms_deltaHz_PostInf | 1 | 5.00% |
| avg_standards_20to50hz_post_stim_post_inf_slope | 1 | 5.00% |
| Peak_raw_251_500ms_Full_spect_PostInf | 1 | 5.00% |
| Pow_251-500ms_thetaHz_PostSubPre | 1 | 5.00% |
| Pow_251-500ms_thetaHz_PostInf | 1 | 5.00% |

Sup**plementary Table S4.4. selected EEG biomarkers at alpha = 0.5 (all drug comparisons combined)**

| EEG biomarker | # of selections | % of selections |
|---|---|---|
| avg_standards_20to50hz_post_stim_post_sub_pre_slope | 16 | 80.00% |
| Pow_1-500ms_thetaHz_PostSubPre | 2 | 10.00% |
| Pow_251-500ms_thetaHz_PostSubPre | 1 | 5.00% |
| Pow_-250--1ms_betaHz_PostSubPre | 1 | 5.00% |
| Pow_1-250ms_betaHz_PostSubPre | 1 | 5.00% |
| avg_standards_20to50hz_pre_stim_post_sub_pre_slope | 1 | 5.00% |

**Supplementary Table S5. selected EEG biomarkers (all drug comparisons and alpha values combined)**

| EEG biomarker | # of selections | % of selections |
|---|---|---|
| avg_standards_20to50hz_post_stim_post_sub_pre_slope | 71 | 88.75% |
| Pow_1-250ms_betaHz_PostSubPre | 43 | 53.75% |
| avg_standards_delta_pre_stim_post_sub_pre_peak_pw_0s | 27 | 33.75% |
| Pow_1-500ms_thetaHz_PostSubPre | 23 | 28.75% |
| Pow_-250--1ms_betaHz_PostSubPre | 20 | 25.00% |
| Pow_1-500ms_betaHz_PostSubPre | 17 | 21.25% |
| EvokedPow_251_500ms_Full_spect_PostInf | 16 | 20.00% |
| avg_standards_20to50hz_pre_stim_post_sub_pre_slope | 9 | 11.25% |
| Pow_251-500ms_thetaHz_PostSubPre | 9 | 11.25% |
| avg_standards_20to50hz_post_stim_post_inf_slope | 9 | 11.25% |
| LZC_Sig_251-500ms_fullspectHz_PostSubPre | 8 | 10.00% |
| Peak_raw_251_500ms_Full_spect_PostSubPre | 7 | 8.75% |
| Pow_251-500ms_deltaHz_PostInf | 7 | 8.75% |
| Pow_1-500ms_alphaHz_PostSubPre | 6 | 7.50% |
| Peak_raw_251_500ms_Full_spect_PostInf | 6 | 7.50% |
| SNR_rms_raw_251_500ms_Full_spect_PostInf | 6 | 7.50% |
| avg_standards_20to50hz_pre_stim_post_inf_slope | 4 | 5.00% |
| Pow_251-500ms_thetaHz_PostInf | 4 | 5.00% |
| Pow_251-500ms_deltaHz_PostSubPre | 4 | 5.00% |
| Pow_1-250ms_thetaHz_PostSubPre | 4 | 5.00% |
| avg_standards_delta_post_stim_post_inf_peak_pw_1s | 4 | 5.00% |
| avg_standards_beta_post_stim_post_sub_pre_peak_pw_1s | 4 | 5.00% |
| avg_standards_alpha_post_stim_post_sub_pre_peak_cf_0s | 3 | 3.75% |
| avg_standards_alpha_post_stim_post_sub_pre_peak_pw_1s | 3 | 3.75% |
| avg_standards_alpha_post_stim_post_inf_peak_cf_0s | 3 | 3.75% |
| avg_standards_beta_pre_stim_post_sub_pre_peak_bw_0s | 3 | 3.75% |
| avg_standards_beta_post_stim_post_inf_peak_bw_1s | 3 | 3.75% |
| avg_standards_delta_pre_stim_post_inf_peak_pw_0s | 3 | 3.75% |
| Pow_-250--1ms_alphaHz_PostSubPre | 3 | 3.75% |
| LZC_Sig_251-500ms_fullspectHz_PostInf | 3 | 3.75% |
| avg_standards_theta_pre_stim_post_sub_pre_peak_cf_0s | 3 | 3.75% |
| Pow_1-500ms_deltaHz_PostInf | 2 | 2.50% |
| SNR_rms_raw_251_500ms_Full_spect_PostSubPre | 2 | 2.50% |
| Pow_1-250ms_thetaHz_PostInf | 2 | 2.50% |
| avg_standards_delta_pre_stim_post_sub_pre_peak_cf_0s | 2 | 2.50% |
| Pow_1-250ms_betaHz_PostInf | 2 | 2.50% |
| avg_standards_beta_pre_stim_post_inf_peak_cf_0s | 2 | 2.50% |
| Pow_251-500ms_betaHz_PostSubPre | 2 | 2.50% |
| Pow_-250--1ms_thetaHz_PostSubPre | 2 | 2.50% |
| avg_standards_delta_post_stim_post_sub_pre_peak_cf_1s | 2 | 2.50% |
| avg_standards_gammato50_pre_stim_post_inf_peak_pw_1s | 1 | 1.25% |
| avg_standards_beta_post_stim_post_inf_peak_cf_1s | 1 | 1.25% |
| avg_standards_beta_post_stim_post_inf_peak_pw_0s | 1 | 1.25% |
| avg_standards_theta_pre_stim_post_inf_peak_bw_0s | 1 | 1.25% |
| avg_standards_delta_pre_stim_post_inf_peak_cf_0s | 1 | 1.25% |
| avg_standards_alpha_post_stim_post_sub_pre_peak_pw_0s | 1 | 1.25% |

| | | |
|---|---|---|
| Peak_raw_1_250ms_Full_spect_PostInf | 1 | 1.25% |
| Pow_1-250ms_alphaHz_PostSubPre | 1 | 1.25% |
| avg_standards_delta_post_stim_post_sub_pre_peak_pw_0s | 1 | 1.25% |
| EvokedPow_251_500ms_Full_spect_PostSubPre | 1 | 1.25% |
| avg_standards_gammato50_post_stim_post_inf_peak_pw_0s | 1 | 1.25% |
| avg_standards_beta_post_stim_post_sub_pre_peak_bw_0s | 1 | 1.25% |
| Pow_-250--1ms_deltaHz_PostInf | 1 | 1.25% |
| avg_standards_gammato50_pre_stim_post_sub_pre_peak_pw_0s | 1 | 1.25% |
| avg_standards_theta_pre_stim_post_sub_pre_peak_bw_0s | 1 | 1.25% |
| avg_standards_gammato50_post_stim_post_sub_pre_peak_cf_1s | 1 | 1.25% |
| avg_standards_beta_pre_stim_post_sub_pre_peak_bw_1s | 1 | 1.25% |
| avg_standards_alpha_post_stim_post_sub_pre_peak_cf_1s | 1 | 1.25% |
| avg_standards_beta_pre_stim_post_sub_pre_peak_cf_0s | 1 | 1.25% |
| Pow_-250--1ms_gamma2Hz_PostSubPre | 1 | 1.25% |
| Pow_-250--1ms_betaHz_PostInf | 1 | 1.25% |
| avg_standards_gammato50_post_stim_post_sub_pre_peak_pw_0s | 1 | 1.25% |
| avg_standards_theta_pre_stim_post_sub_pre_peak_pw_0s | 1 | 1.25% |
| avg_standards_gammato50_pre_stim_post_sub_pre_peak_pw_1s | 1 | 1.25% |

#### Supplementary Tables S6-S9: selected EEG biomarker categories

##### S6. Supplementary Tables: selected EEG biomarker categories for each drug comparison (all alpha values combined)

**Supplementary Table S6.1. Selected EEG biomarker categories for the ketamine, thiopental, and placebo comparison (all alpha values combined)**

| Biomarker category | # of selections | % of selections |
|---|---|---|
| PSD exponent (aperiodic) | 20 | 100.00% |
| Beta power (total) | 13 | 65.00% |
| Theta power (total) | 12 | 60.00% |
| Delta power 1st peak (periodic) | 10 | 50.00% |
| P300 evoked power | 9 | 45.00% |
| P300 peak amplitude | 2 | 10.00% |
| Alpha power (total) | 2 | 10.00% |
| Theta power 1st peak (periodic) | 1 | 5.00% |
| Delta power (total) | 1 | 5.00% |
| Beta power 2nd peak (periodic) | 1 | 5.00% |
| Alpha center frequency 1st peak (periodic) | 1 | 5.00% |
| Gamma 2 power (total) | 1 | 5.00% |
| Alpha power 2nd peak (periodic) | 1 | 5.00% |
| Lempel-Ziv complexity | 1 | 5.00% |
| Beta center frequency 1st peak (periodic) | 1 | 5.00% |
| Gamma power 1st peak (periodic) | 1 | 5.00% |
| P300 signal-to-noise ratio | 1 | 5.00% |

**Supplementary Table S6.2. Selected EEG biomarker categories for the ketamine and placebo comparison (all alpha values combined)**

| Biomarker category | # of selections | % of selections |
|---|---|---|
| PSD exponent (aperiodic) | 19 | 95.00% |
| Beta power (total) | 13 | 65.00% |
| Delta power 1st peak (periodic) | 12 | 60.00% |
| P300 peak amplitude | 7 | 35.00% |
| Delta power (total) | 7 | 35.00% |
| Delta power 2nd peak (periodic) | 4 | 20.00% |
| Alpha power (total) | 4 | 20.00% |
| Beta bandwidth 2nd peak (periodic) | 4 | 20.00% |
| Alpha center frequency 1st peak (periodic) | 4 | 20.00% |
| Beta bandwidth 1st peak (periodic) | 3 | 15.00% |
| Beta power 2nd peak (periodic) | 3 | 15.00% |
| Alpha power 2nd peak (periodic) | 2 | 10.00% |
| Gamma power 1st peak (periodic) | 2 | 10.00% |
| Delta center frequency 2nd peak (periodic) | 2 | 10.00% |
| P300 evoked power | 2 | 10.00% |
| Beta center frequency 1st peak (periodic) | 2 | 10.00% |
| Gamma power 2nd peak (periodic) | 1 | 5.00% |
| Alpha center frequency 2nd peak (periodic) | 1 | 5.00% |
| Theta power (total) | 1 | 5.00% |
| Delta center frequency 1st (periodic) | 1 | 5.00% |
| Gamma center frequency 2nd (periodic) | 1 | 5.00% |
| Theta bandwidth 1st peak (periodic) | 1 | 5.00% |

**Supplementary Table S6.3. Selected EEG biomarker categories for the thiopental and placebo comparison (all alpha values combined)**

| Biomarker category | # of selections | % of selections |
|---|---|---|
| Theta power (total) | 18 | 90.00% |
| Beta power (total) | 15 | 75.00% |
| PSD exponent (aperiodic) | 13 | 65.00% |
| Lempel-Ziv complexity | 7 | 35.00% |
| Delta power 1st peak (periodic) | 6 | 30.00% |
| P300 evoked power | 6 | 30.00% |
| P300 signal-to-noise ratio | 5 | 25.00% |
| P300 peak amplitude | 4 | 20.00% |
| Delta power (total) | 3 | 15.00% |
| Alpha power (total) | 3 | 15.00% |
| Theta center frequency 1st peak (periodic) | 3 | 15.00% |
| Delta center frequency 1st peak (periodic) | 2 | 10.00% |
| N100 peak amplitude | 1 | 5.00% |
| Theta bandwidth 1st peak (periodic) | 1 | 5.00% |
| Beta center frequency 2nd peak (periodic) | 1 | 5.00% |
| Beta power 1st peak (periodic) | 1 | 5.00% |
| Alpha power 1st peak (periodic) | 1 | 5.00% |
| Gamma power 2nd peak (periodic) | 1 | 5.00% |

**Supplementary Table S6.4. Selected EEG biomarker categories for the ketamine and thiopental comparison (all alpha values combined)**

| Biomarker category | # of selections | % of selections |
|---|---|---|
| PSD exponent (aperiodic) | 20 | 100.00% |
| Beta power (total) | 10 | 50.00% |
| Theta power (total) | 1 | 5.00% |

#### S7. Supplementary Tables: selected EEG biomarker categories for each drug comparison across alpha values

**Supplementary Table S7.1. Selected EEG biomarker categories for the ketamine, thiopental, and placebo comparison at alpha = 0.01**

| Biomarker category | # of selections | % of selections |
|---|---|---|
| Beta power (total) | 5 | 100.00% |
| Theta power (total) | 5 | 100.00% |
| PSD exponent (aperiodic) | 5 | 100.00% |
| P300 evoked power | 4 | 80.00% |
| Delta power 1st peak (periodic) | 4 | 80.00% |
| Alpha power (total) | 2 | 40.00% |
| Gamma power 1st peak (periodic) | 1 | 20.00% |
| Gamma 2 power (total) | 1 | 20.00% |
| Beta center frequency 1st peak (periodic) | 1 | 20.00% |
| P300 peak amplitude | 1 | 20.00% |
| Lempel-Ziv complexity | 1 | 20.00% |
| Alpha power 2nd peak (periodic) | 1 | 20.00% |
| Theta power 1st peak (periodic) | 1 | 20.00% |
| Alpha center frequency 1st peak (periodic) | 1 | 20.00% |
| Beta power 2nd peak (periodic) | 1 | 20.00% |

**Supplementary Table S7.2. Selected EEG biomarker categories for the ketamine, thiopental, and placebo comparison at alpha = 0.1**

| Biomarker category | # of selections | % of selections |
|---|---|---|
| PSD exponent (aperiodic) | 5 | 100.00% |
| Beta power (total) | 5 | 100.00% |
| Theta power (total) | 5 | 100.00% |
| Delta power 1st peak (periodic) | 4 | 80.00% |
| P300 evoked power | 3 | 60.00% |
| P300 peak amplitude | 1 | 20.00% |
| Delta power (total) | 1 | 20.00% |
| P300 signal-to-noise ratio | 1 | 20.00% |

**Supplementary Table S7.3. Selected EEG biomarker categories for the ketamine, thiopental, and placebo comparison at alpha = 0.2**

| Biomarker category | # of selections | % of selections |
|---|---|---|
| PSD exponent (aperiodic) | 5 | 100.00% |
| Beta power (total) | 3 | 60.00% |
| Theta power (total) | 2 | 40.00% |
| P300 evoked power | 2 | 40.00% |
| Delta power 1st peak (periodic) | 2 | 40.00% |

**Supplementary Table S7.4. Selected EEG biomarker categories for the ketamine, thiopental, and placebo comparison at alpha = 0.5**

| Biomarker category | # of selections | % of selections |
|---|---|---|
| PSD exponent (aperiodic) | 5 | 100.00% |

**Supplementary Table S7.5. Selected EEG biomarker categories for the ketamine and placebo comparison at alpha = 0.01**

| Biomarker category | # of selections | % of selections |
|---|---|---|
| Beta power (total) | 5 | 100.00% |
| Delta power 1st peak (periodic) | 5 | 100.00% |
| PSD exponent (aperiodic) | 5 | 100.00% |
| Delta power 2nd peak (periodic) | 4 | 80.00% |
| Alpha power (total) | 4 | 80.00% |
| Delta power (total) | 4 | 80.00% |
| Alpha center frequency 1st peak (periodic) | 3 | 60.00% |
| Beta bandwidth 2nd peak (periodic) | 3 | 60.00% |
| P300 peak amplitude | 3 | 60.00% |
| Beta bandwidth 1st peak (periodic) | 2 | 40.00% |
| P300 evoked power | 2 | 40.00% |
| Gamma power 1st peak (periodic) | 2 | 40.00% |
| Alpha power 2nd peak (periodic) | 2 | 40.00% |
| Beta center frequency 1st peak (periodic) | 2 | 40.00% |
| Delta center frequency 2nd peak (periodic) | 2 | 40.00% |
| Beta power 2nd peak (periodic) | 2 | 40.00% |
| Gamma power 2nd peak (periodic) | 1 | 20.00% |
| Alpha center frequency 2nd peak (periodic) | 1 | 20.00% |
| Theta power (total) | 1 | 20.00% |
| Delta center frequency 1st peak (periodic) | 1 | 20.00% |
| Gamma center frequency 2nd peak (periodic) | 1 | 20.00% |
| Theta bandwidth 1st peak (periodic) | 1 | 20.00% |

**Supplementary Table S7.6 selected EEG biomarker categories for the ketamine and placebo comparison at alpha = 0.1**

| Biomarker category | # of selections | % of selections |
|---|---|---|
| PSD exponent (aperiodic) | 5 | 100.00% |
| Beta power (total) | 4 | 80.00% |
| Delta power 1st peak (periodic) | 4 | 80.00% |
| P300 peak amplitude | 2 | 40.00% |
| Delta power (total) | 2 | 40.00% |
| Beta power 2nd peak (periodic) | 1 | 20.00% |
| Alpha center frequency 1st peak (periodic) | 1 | 20.00% |
| Beta bandwidth 1st peak (periodic) | 1 | 20.00% |
| Beta bandwidth 2nd peak (periodic) | 1 | 20.00% |

**Supplementary Table S7.7. Selected EEG biomarker categories for the ketamine and placebo comparison at alpha = 0.2**

| Biomarker category | # of selections | % of selections |
|---|---|---|
| Beta power (total) | 4 | 80.00% |
| PSD exponent (aperiodic) | 4 | 80.00% |
| Delta power 1st peak (periodic) | 3 | 60.00% |
| P300 peak amplitude | 2 | 40.00% |
| Delta power (total) | 1 | 20.00% |

**Supplementary Table S7.8. Selected EEG biomarker categories for the ketamine and placebo comparison at alpha = 0.5**

| Biomarker category | # of selections | % of selections |
|---|---|---|
| PSD exponent (aperiodic) | 5 | 100.00% |

**Supplementary Table S7.9. Selected EEG biomarker categories for the thiopental and placebo comparison at alpha = 0.01**

| Biomarker category | # of selections | % of selections |
|---|---|---|
| Lempel-Ziv complexity | 5 | 100.00% |
| P300 signal-to-noise ratio | 5 | 100.00% |
| Delta power 1st peak (periodic) | 5 | 100.00% |
| Theta power (total) | 5 | 100.00% |
| PSD exponent (aperiodic) | 5 | 100.00% |
| Beta power (total) | 5 | 100.00% |
| Alpha power (total) | 3 | 60.00% |
| P300 peak amplitude | 2 | 40.00% |
| P300 evoked power | 2 | 40.00% |
| Delta center frequency 1st peak (periodic) | 2 | 40.00% |
| Theta center frequency 1st peak (periodic) | 2 | 40.00% |
| Delta power (total) | 2 | 40.00% |
| N100 peak amplitude | 1 | 20.00% |
| Theta bandwidth 1st peak (periodic) | 1 | 20.00% |
| Beta center frequency 2nd peak (periodic) | 1 | 20.00% |
| Beta power 1st peak (periodic) | 1 | 20.00% |
| Alpha power 1st peak (periodic) | 1 | 20.00% |

**Supplementary Table S7.10. Selected EEG biomarker categories for the thiopental and placebo comparison at alpha = 0.1**

| Biomarker category | # of selections | % of selections |
|---|---|---|
| PSD exponent (aperiodic) | 5 | 100.00% |
| Theta power (total) | 5 | 100.00% |
| Beta power (total) | 4 | 80.00% |
| P300 evoked power | 3 | 60.00% |
| Lempel-Ziv complexity | 2 | 40.00% |
| Delta power 1st peak (periodic) | 1 | 20.00% |
| Gamma power 2nd peak (periodic) | 1 | 20.00% |
| Theta center frequency 1st peak (periodic) | 1 | 20.00% |
| Delta power (total) | 1 | 20.00% |
| P300 peak amplitude | 1 | 20.00% |

**Supplementary Table S7.11. Selected EEG biomarker categories for the thiopental and placebo comparison at alpha = 0.2**

| Biomarker category | # of selections | % of selections |
|---|---|---|
| Theta power (total) | 5 | 100.00% |
| Beta power (total) | 4 | 80.00% |
| PSD exponent (aperiodic) | 1 | 20.00% |
| P300 evoked power | 1 | 20.00% |
| P300 peak amplitude | 1 | 20.00% |

**Supplementary Table S7.12. Selected EEG biomarker categories for the thiopental and placebo comparison at alpha = 0.5**

| Biomarker category | # of selections | % of selections |
|---|---|---|
| Theta power (total) | 3 | 60.00% |
| Beta power (total) | 2 | 40.00% |
| PSD exponent (aperiodic) | 2 | 40.00% |

**Supplementary Table S7.13. Selected EEG biomarker categories for the ketamine and thiopental comparison at alpha = 0.01**

| Biomarker category | # of selections | % of selections |
|---|---|---|
| PSD exponent (aperiodic) | 5 | 100.00% |
| Beta power (total) | 5 | 100.00% |
| Theta power (total) | 1 | 20.00% |

**Supplementary Table S7.14. Selected EEG biomarker categories for the ketamine and thiopental comparison at alpha = 0.1**

| Biomarker category | # of selections | % of selections |
|---|---|---|
| PSD exponent (aperiodic) | 5 | 100.00% |
| Beta power (total) | 5 | 100.00% |

**Supplementary Table S7.15. Selected EEG biomarker categories for the ketamine and thiopental comparison at alpha = 0.2**

| Biomarker category | # of selections | % of selections |
|---|---|---|
| PSD exponent (aperiodic) | 5 | 100.00% |

**Supplementary Table S7.16. Selected EEG biomarker categories for the ketamine and thiopental comparison at alpha = 0.5**

| Biomarker category | # of selections | % of selections |
|---|---|---|
| PSD exponent (aperiodic) | 5 | 100.00% |

**S8. Supplementary Tables: selected EEG biomarker categories across alpha values (all drug comparisons combined)**

**Supplementary Table s8.1. Selected EEG biomarker categories at alpha = 0.01 (all drug comparisons combined)**

| Biomarker category | # of selections | % of selections |
|---|---|---|
| Beta power (total) | 20 | 100.00% |
| PSD exponent (aperiodic) | 20 | 100.00% |
| Delta power 1st peak (periodic) | 14 | 70.00% |
| Theta power (total) | 12 | 60.00% |
| Alpha power (total) | 9 | 45.00% |
| P300 evoked power | 8 | 40.00% |
| P300 peak amplitude | 7 | 35.00% |
| Lempel-Ziv complexity | 6 | 30.00% |
| Delta power (total) | 6 | 30.00% |
| P300 signal-to-noise ratio | 5 | 25.00% |
| Alpha center frequency 1st (periodic) | 4 | 20.00% |
| Delta power 2nd peak (periodic) | 4 | 20.00% |
| Delta center frequency 1st peak (periodic) | 3 | 15.00% |
| Beta bandwidth 2nd peak (periodic) | 3 | 15.00% |
| Beta power 2nd peak (periodic) | 3 | 15.00% |
| Alpha power 2nd peak (periodic) | 3 | 15.00% |
| Beta center frequency 1st peak (periodic) | 3 | 15.00% |
| Gamma power 1st peak (periodic) | 3 | 15.00% |
| Beta bandwidth 1st peak (periodic) | 2 | 10.00% |
| Delta center frequency 2nd peak (periodic) | 2 | 10.00% |
| Theta bandwidth 1st peak (periodic) | 2 | 10.00% |
| Theta center frequency 1st peak (periodic) | 2 | 10.00% |
| N100 peak amplitude | 1 | 5.00% |
| Gamma 2 power (total) | 1 | 5.00% |
| Theta power 1st peak (periodic) | 1 | 5.00% |
| Gamma power 2nd peak (periodic) | 1 | 5.00% |
| Alpha center frequency 2nd peak (periodic) | 1 | 5.00% |
| Gamma center frequency 2nd peak (periodic) | 1 | 5.00% |
| Beta center frequency 2nd peak (periodic) | 1 | 5.00% |
| Beta power 1st peak (periodic) | 1 | 5.00% |
| Alpha power 1st peak (periodic) | 1 | 5.00% |

**Supplementary Table s8.2. Selected EEG biomarker categories at alpha = 0.1 (all drug comparisons combined)**

| Biomarker category | # of selections | % of selections |
|---|---|---|
| PSD exponent (aperiodic) | 20 | 100.00% |
| Beta power (total) | 18 | 90.00% |
| Theta power (total) | 10 | 50.00% |
| Delta power (periodic) | 9 | 45.00% |
| P300 evoked power | 6 | 30.00% |
| P300 peak amplitude | 4 | 20.00% |
| Delta power (total) | 4 | 20.00% |
| Lempel-Ziv complexity | 2 | 10.00% |
| P300 signal-to-noise ratio | 1 | 5.00% |
| Beta power 2$^{nd}$ peak (periodic) | 1 | 5.00% |
| Alpha center frequency 1$^{st}$ peak (periodic) | 1 | 5.00% |
| Beta bandwidth 1$^{st}$ peak (periodic) | 1 | 5.00% |
| Beta bandwidth 2$^{nd}$ peak (periodic) | 1 | 5.00% |
| Gamma power 2$^{nd}$ peak (periodic) | 1 | 5.00% |
| Theta center frequency 1$^{st}$ peak (periodic) | 1 | 5.00% |

**Supplementary Table s8.3. Selected EEG biomarker categories at alpha = 0.2 (all drug comparisons combined)**

| Biomarker category | # of selections | % of selections |
|---|---|---|
| PSD exponent (aperiodic) | 15 | 75.00% |
| Beta power (total) | 11 | 55.00% |
| Theta power (total) | 7 | 35.00% |
| Delta power 1$^{st}$ peak (periodic) | 5 | 25.00% |
| P300 evoked power | 3 | 15.00% |
| P300 peak amplitude | 3 | 15.00% |
| Delta power (total) | 1 | 5.00% |

**Supplementary Table s8.4. Selected EEG biomarker categories at alpha = 0.5 (all drug comparisons combined)**

| Biomarker category | # of selections | % of selections |
|---|---|---|
| PSD exponent (aperiodic) | 17 | 85.00% |
| Theta power (total) | 3 | 15.00% |
| Beta power (total) | 2 | 10.00% |

**Supplementary Table s9. Selected EEG biomarker categories (all drug comparisons and alpha values combined)**

| Biomarker category | # of selections | % of selections |
|---|---|---|
| PSD exponent (aperiodic) | 72 | 90.00% |
| Beta power (total) | 51 | 63.75% |
| Theta power (total) | 32 | 40.00% |
| Delta power 1st peak (periodic) | 28 | 35.00% |
| P300 evoked power | 17 | 21.25% |
| P300 peak amplitude | 13 | 17.50% |
| Delta power (total) | 11 | 13.75% |
| Alpha power (total) | 9 | 11.25% |
| Lempel-Ziv complexity | 8 | 10.00% |
| P300 signal-to-noise ratio | 6 | 7.50% |
| Alpha center frequency 1st peak (periodic) | 5 | 6.25% |
| Beta power 2nd peak (periodic) | 4 | 5.00% |
| Delta power 2nd peak (periodic) | 4 | 5.00% |
| Beta bandwidth 2nd peak (periodic) | 4 | 5.00% |
| Theta center frequency 1st peak (periodic) | 3 | 3.75% |
| Beta bandwidth 1st peak (periodic) | 3 | 3.75% |
| Delta center frequency 1st peak (periodic) | 3 | 3.75% |
| Beta center frequency 1st peak (periodic) | 3 | 3.75% |
| Alpha power 2nd peak (periodic) | 3 | 3.75% |
| Gamma power 1st peak (periodic) | 3 | 3.75% |
| Gamma power 2nd peak (periodic) | 2 | 2.50% |
| Delta center frequency 2nd peak (periodic) | 2 | 2.50% |
| Theta bandwidth 1st peak (periodic) | 2 | 2.50% |
| N100 peak amplitude | 1 | 1.25% |
| Gamma 2 power (total) | 1 | 1.25% |
| Theta power 1st peak (periodic) | 1 | 1.25% |
| Alpha center frequency 2nd peak (periodic) | 1 | 1.25% |
| Gamma center frequency 2nd peak (periodic) | 1 | 1.25% |
| Beta center frequency 2nd peak (periodic) | 1 | 1.25% |
| Beta power 1st peak (periodic) | 1 | 1.25% |
| Alpha power 1st peak (periodic) | 1 | 1.25% |

#### Supplementary Tables S10: Classification metrics

Each table reports the mean and standard deviation (in parentheses) of the classification metrics for each drug comparison. The metrics were calculated across the test splits of a five-fold cross-validation procedure independently for each value of the selection size restriction parameter alpha. The second column shows the mean number of selected EEG biomarkers and standard deviation (in parentheses) across the training splits. Accuracy refers to the percentage of correct drug classifications out of the total number of cases; precision refers to the percentage of correct classifications for a given drug out of all the cases predicted as that drug; and sensitivity refers to the percentage of correct classifications for a given drug out of all cases that received that drug.

**Supplementary Table S10.1. Metrics for the 3-class placebo, ketamine, and thiopental classification**

| Alpha | # selected biomarkers | Accuracy | Precision | | | Sensitivity | | |
|---|---|---|---|---|---|---|---|---|
| | | | Placebo | Ketamine | Thiopental | Placebo | Ketamine | Thiopental |
| All features | 187.00<br>(0.00) | 90.50<br>(2.90) | 88.51<br>(9.92) | 91.62<br>(5.94) | 100.00<br>(0.00) | 91.56<br>(5.18) | 92.46<br>(5.32) | 89.06<br>(7.80) |
| 0.01 | 9.80<br>(3.06) | 89.01<br>(2.01) | 83.93<br>(8.73) | 93.09<br>(6.30) | 97.13<br>(3.53) | 85.98<br>(7.59) | 85.81<br>(6.53) | 95.38<br>(6.15) |
| 0.1 | 5.40<br>(1.02) | 88.09<br>(3.91) | 86.13<br>(5.89) | 92.50<br>(7.29) | 89.60<br>(5.97) | 83.20<br>(9.33) | 90.00<br>(5.96) | 95.49<br>(3.68) |
| 0.2 | 2.80<br>(0.40) | 78.17<br>(5.86) | 70.03<br>(6.27) | 83.40<br>(10.64) | 86.51<br>(8.36) | 71.50<br>(13.61) | 83.16<br>(7.57) | 80.74<br>(7.19) |
| 0.5 | 1.00<br>(0.00) | 67.85<br>(11.18) | 61.22<br>(15.45) | 80.31<br>(12.75) | 65.70<br>(14.98) | 58.32<br>(15.93) | 77.21<br>(13.62) | 67.70<br>(14.27) |

**Supplementary Table S10.2. Metrics for the 2-class placebo and ketamine classification**

| Alpha | # selected biomarkers | Accuracy | Precision | | Sensitivity | |
|---|---|---|---|---|---|---|
| | | | Placebo | Ketamine | Placebo | Ketamine |
| All features | 187.00<br>(0.00) | 94.05<br>(2.27) | 90.90<br>(3.83) | 92.39<br>(4.48) | 91.83<br>(4.78) | 92.15<br>(2.34) |
| 0.01 | 15.40<br>(3.44) | 90.69<br>(4.60) | 91.46<br>(2.96) | 90.18<br>(6.87) | 85.08<br>(10.83) | 90.90<br>(4.84) |
| 0.1 | 4.80<br>(1.17) | 87.13<br>(3.72) | 87.50<br>(9.45) | 87.04<br>(3.72) | 86.33<br>(9.09) | 85.38<br>(5.43) |
| 0.2 | 3.00<br>(0.63) | 83.16<br>(5.69) | 82.45<br>(4.17) | 83.22<br>(12.29) | 80.83<br>(13.78) | 82.71<br>(5.77) |
| 0.5 | 1.00<br>(0.00) | 77.95<br>(12.90) | 78.81<br>(14.95) | 79.38<br>(13.02) | 77.50<br>(16.21) | 79.07<br>(15.70) |

**Supplementary Table S10.3. Metrics for the 2-class placebo and thiopental classification**

| Alpha | # selected biomarkers | Accuracy | Precision | | Sensitivity | |
|---|---|---|---|---|---|---|
| | | | Placebo | Thiopental | Placebo | Thiopental |
| All features | 187.00<br>(0.00) | 92.49<br>(3.37) | 93.39<br>(8.12) | 97.33<br>(5.33) | 97.03<br>(3.64) | 88.98<br>(8.21) |
| 0.01 | 16.00<br>(1.90) | 94.80<br>(3.90) | 91.09<br>(4.41) | 94.85<br>(4.35) | 97.13<br>(3.53) | 88.16<br>(4.71) |
| 0.1 | 5.00<br>(1.26) | 90.95<br>(2.24) | 89.57<br>(4.55) | 90.13<br>(8.26) | 94.45<br>(2.82) | 91.98<br>(5.33) |
| 0.2 | 2.60<br>(0.49) | 84.93<br>(4.31) | 86.14<br>(4.25) | 83.30<br>(4.86) | 85.94<br>(5.36) | 82.83<br>(7.35) |
| 0.5 | 1.40<br>(0.49) | 70.71<br>(9.48) | 75.19<br>(9.53) | 65.87<br>(9.98) | 69.62<br>(10.36) | 71.80<br>(16.90) |

**Supplementary Table S10.4. Metrics for the 2-class ketamine and thiopental classification**

| Alpha | # selected biomarkers | Accuracy | Precision | | Sensitivity | |
|---|---|---|---|---|---|---|
| | | | Ketamine | Thiopental | Ketamine | Thiopental |
| All features | 178.00<br>(0.00) | 97.00<br>(1.50) | 94.96<br>(4.71) | 98.57<br>(2.86) | 98.57<br>(2.86) | 95.03<br>(4.17) |
| 0.01 | 3.20<br>(1.17) | 97.08<br>(1.48) | 97.49<br>(3.08) | 96.85<br>(3.94) | 98.57<br>(2.86) | 98.57<br>(2.86) |
| 0.1 | 2.00<br>(0.00) | 98.62<br>(1.69) | 100.00<br>(0.00) | 98.67<br>(2.67) | 98.57<br>(2.86) | 100.00<br>(0.00) |
| 0.2 | 1.00<br>(0.00) | 94.91<br>(2.82) | 94.82<br>(4.96) | 94.83<br>(4.26) | 96.06<br>(3.24) | 94.18<br>(5.38) |
| 0.5 | 1.00<br>(0.00) | 94.91<br>(2.82) | 94.82<br>(4.96) | 94.83<br>(4.26) | 96.06<br>(3.24) | 94.18<br>(5.38) |
